# Genome-wide neighbor effects predict genotype pairs that reduce herbivory in mixed planting

**DOI:** 10.1101/2023.05.19.541411

**Authors:** Yasuhiro Sato, Rie Shimizu-Inatsugi, Kazuya Takeda, Bernhard Schmid, Atsushi J. Nagano, Kentaro K. Shimizu

**Affiliations:** Department of Evolutionary Biology and Environmental Studies, University of Zurich, Winterthurerstrasse 190, 8057 Zurich, Switzerland; Research Institute for Food and Agriculture, Ryukoku University, Yokotani 1-5, Seta Oe-cho, Otsu, Shiga 520-2194, Japan; Department of Geography, University of Zurich, Winterthurerstrasse 190, 8057 Zurich, Switzerland; Faculty of Agriculture Ryukoku University, Yokotani 1-5, Seta Oe-cho, Otsu, Shiga 520-2194, Japan; Institute for Advanced Biosciences, Keio University, 403-1 Nipponkoku, Daihouji, Tsuruoka, Yamagata, 997-0017, Japan; Kihara Institute for Biological Research, Yokohama City University, Maioka 641-12, Totsuka-ward, Yokohama 244-0813, Japan

**Author notes:** Co-corresponding authors (Y.S.); (A.J.N); (K.K.S.).

**Keywords:** Associational resistance, Genetic diversity, Plant-herbivore interaction

## Abstract

Genetically diverse populations can increase plant resistance to natural enemies. Yet, beneficial genotype pairs remain elusive due to the occurrence of both positive and negative effects of mixed planting on plant resistance, called associational resistance and susceptibility. We used genome-wide polymorphisms of the plant species *Arabidopsis thaliana* to identify genotype pairs that enhance associational resistance to herbivory. By quantifying neighbor interactions among 199 genotypes grown in a randomized block design, we predicted that 823 of the 19,701 candidate pairs could reduce herbivory through associational resistance. We planted such pairs with predicted associational resistance in mixtures and monocultures and found a significant reduction in herbivore damage in the mixtures. Our study highlights the potential application to assemble genotype mixtures with positive biodiversity effects.

## Main Text

Genetic diversity is increasingly recognized as a critical facet of biodiversity (*1, 2*) that should be conserved as a provider of various ecosystem services (*3*) as well as a source of evolution (*2, 4*). In terrestrial ecosystems, for example, plant genotypic diversity can increase plant resistance to natural enemies as the number of plant genotypes in a contiguous group of plants, namely a stand, increases (*5–7*). However, such a stand of multiple plant genotypes does not always result in positive outcomes (*8–10*). Identifying beneficial pairs from a mixture of genotypes helps us design a desirable mixture and understand the potential mechanisms affecting stand-level properties.

Both positive and negative effects of mixed planting on stand-level resistance to herbivores have been reported in the literature (*7, 11–13*). The underlying mechanisms have been referred to as associational resistance and associational susceptibility, respectively (*11*). Because plants are sessile, such associational resistance and susceptibility are driven by plant-plant interactions among neighboring individuals (*11*). If resistant plants repel herbivores and thereby protect susceptible neighbors, associational resistance occurs rendering a mixture of resistant and susceptible plants less damaged than corresponding monocultures (*14, 15*). In contrast, associational susceptibility leads the mixture to incur more damage if herbivores are attracted to susceptible plants and then spill onto resistant neighbors (*8, 14*). The combined occurrence of associational resistance and susceptibility in a single mixture makes it difficult to distinguish between positively and negatively interacting genotype pairs for anti-herbivore resistance.

Recent studies have used standard genome-wide association studies (GWAS) to dissect the genetic basis underlying beneficial plant-plant interactions (*16, 17*). However, it is difficult to identify the most beneficial pairs among many potential pairs. In this study, we aimed to predict such pairs by combining genome-wide single nucleotide polymorphisms (SNPs) in *Arabidopsis thaliana* (*18, 19*) with a new GWAS method named “Neighbor GWAS” (*20*). Neighbor GWAS adopts a physical model of magnets to estimate locus-wise positive or negative interactions between focal and neighbor individuals over randomized mixtures of many genotypes (*20*) (Fig. 1). We first planted replicated individuals of 199 *A. thaliana* genotypes at two field sites and observed naturally emerging communities of herbivores, which were analyzed as extended phenotypes of the plants in standard GWAS or Neighbor GWAS. We then used Neighbor GWAS as a tool to predict associational resistance or susceptibility out of all possible 19,701 pairs among the 199 genotypes. To test our prediction, we finally planted genotypes of prospective beneficial pairs in mixtures and monocultures.

**Figure 1.**
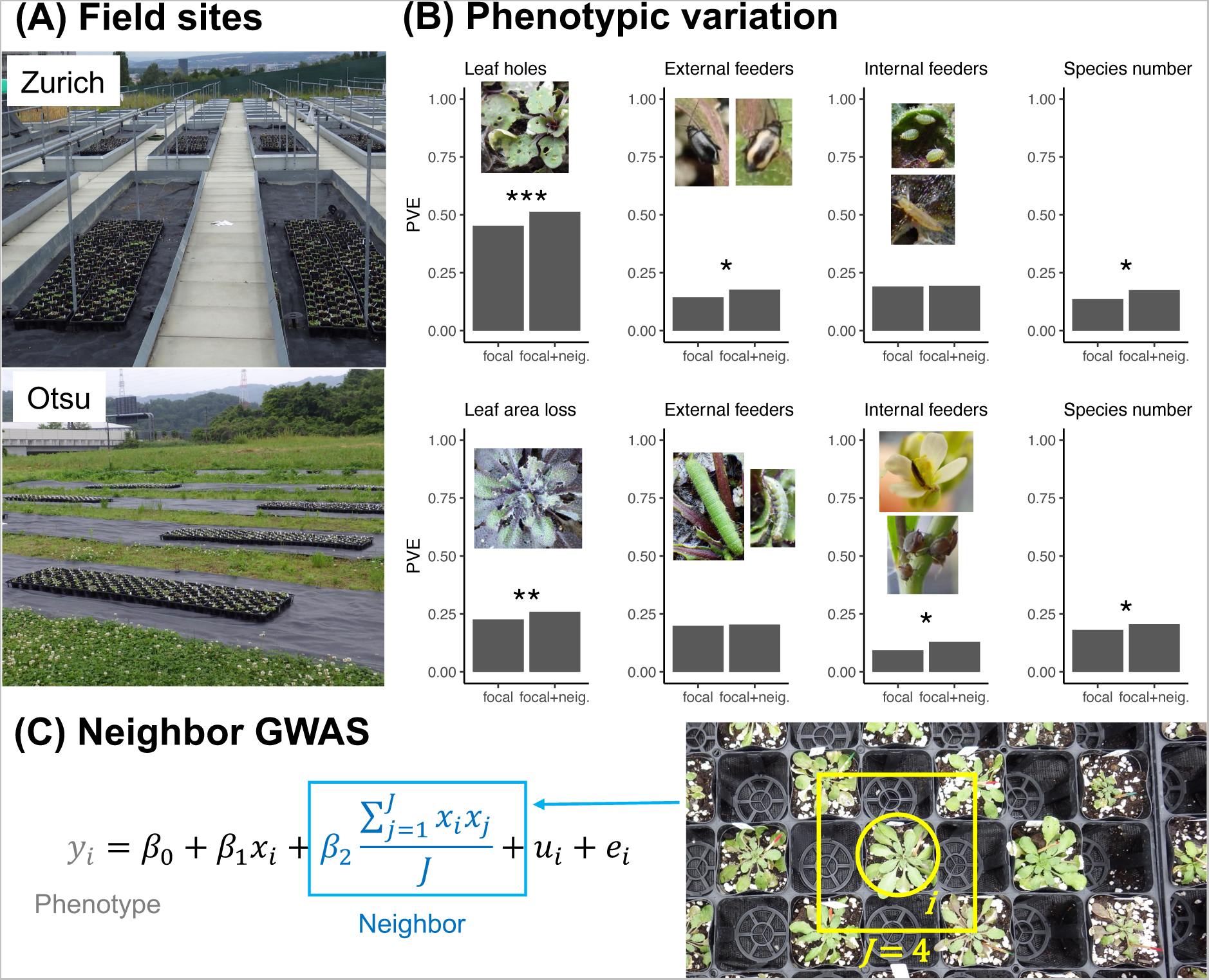
Genetic variation in herbivore damage and community composition on randomized mixtures of *Arabidopsis thaliana* genotypes. (A) 1,600 *A. thaliana* individuals (200 plants × 8 randomized blocks) were planted in the Zurich or Otsu site for two years. Potted plants were arranged in a checkered manner (cf. photograph in C). (B) The proportion of phenotypic variation explained (PVE) by focal genotypes alone (focal) or both focal and neighbor genotypes (focal+neig.). Asterisks highlight the significant contributions of neighbor genotypes over those of focal genotypes: *** *p* < 0.001; ** *p* < 0.01; \**p* < 0.05 (Table S3). (C) Neighbor GWAS model that includes neighbor genotype effects besides focal genotype effects. The term 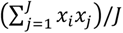 represents the mean allele similarity between the focal (*x_i_*) and neighbor (*x_j_*; j up to *J*) individuals. The coefficients *β*_1_ or *β*_2_ represent single-locus effects of the focal or neighbor genotypes on the phenotype value of the *i*-th focal individual *y_i_*, respectively.

To enable GWAS of herbivore damage, we planted *A. thaliana* genotypes in a randomized block design in two experimental gardens over two years (Table S1; Fig. 1A). This allowed us to monitor the abundance of 18 insect species on nearly 6400 individual plants (≈ 199 genotypes × 8 blocks × 2 sites × 2 years) at a native (Zurich, Switzerland) or exotic (Otsu, Japan) field site (Table S2; Fig. S1). We quantified herbivore damage as the number of leaf holes in Zurich and leaf area loss in Otsu because the major herbivores in Zurich were flea beetles and those in Otsu were diamondback moths or small white butterflies (Fig. 1B; Fig. S1). To specify insect functional groups responsible for herbivore damage, we quantified three extended phenotypes for herbivore communities by counting individuals of external feeders (e.g., beetles in Zurich or caterpillars in Otsu), individuals of internal feeders (aphids and thrips), and all insect species per plant individual (Fig. S2). All four phenotypes exhibited quantitative phenotypic variation among the individual plants (Fig. S2), making them suitable target phenotypes for GWAS.

Before using the Neighbor GWAS, we performed a standard GWAS to examine focal genotype effects on herbivore damage and insect community composition. For all four phenotypes, we found significant heritability among plant genotypes at both the sites (likelihood ratio test, *p* < 0.05: “focal” in Fig. 1B; Fig. S3; Table S3). Regarding the effects of focal genotypes on herbivore damage in Zurich (Fig. S4A; Table S4), we detected a significant SNP in the *GLABRA1* gene. This gene is known to initiate leaf trichome development and thereby prevent herbivory by flea beetles (*21*). Although previous studies reported significant effects of the glucosinolate genes *GS-OH* and *MAM1* on field herbivory (*22*), none of the measured phenotypes showed significant peaks near these glucosinolate genes (Fig. S4A and S5A; Table S4). This was likely because most herbivores observed in this study were specialists (Fig. S1; Table S2) and thus overcame the glucosinolate defense. The results of the standard GWAS agreed with previous evidence for physical defense, whereas the herbivore damage observed in our study was unlikely to be attributable to the known mechanisms of defense by glucosinolates.

To test whether neighbor genotypes contributed to genetic variation in herbivore damage, we applied the Neighbor GWAS method that considered neighbor genotype effects besides the focal genotype effects (Fig. 1C) (*20*). The neighbor genotypes explained a significant fraction of the phenotypic variation in the herbivore damage of focal plants at both sites compared with focal genotype effects alone (“focal+neig.” in Fig. 1B; Fig. S3; Table S3), indicating the importance of neighbor genotypes in shaping herbivore damage. Additionally, we performed Neighbor GWAS of the insect community composition to examine which types of insect herbivores were the most influenced by neighbor genotypes. Flea beetles that could jump between plants were abundant in Zurich (Fig. S1) and its abundance on focal plants was significantly influenced by neighbor genotypes (Fig. 1B; Table S3). In contrast, the contribution of neighbor genotypes to the number of external feeders on focal plants was not significant in Otsu (Fig. 1B; Table S3), where the major external feeders were sedentary caterpillars that did not move between the plants (Fig. S1). Flower thrips that can move between flowering plants were abundant in Otsu (Fig. S1) and the number of internal feeders including this thrip species was significantly influenced by neighbor genotypes (Fig. 1B; Table S3). Reflecting the significant contributions of neighbor genotypes to either external feeders in Zurich or internal feeders in Otsu, neighbor genotypes significantly contributed to the total number of insect species on focal plants at both sites (Fig. 1B; Fig. S3; Table S3). These patterns of herbivore damage and communities suggest that neighbor genotypes are more likely to influence mobile herbivores than sedentary herbivores.

We then asked how many loci underlay the influence of neighbor genotypes on herbivore damage and herbivore communities on focal plants. To attribute the phenotypic variation to each SNP, we mapped the statistical significance of the neighbor genotype effect *β*_2_ throughout the *A. thaliana* genome (Fig. 2A and B). This association mapping did not detect any significant SNPs for any of the four phenotypes at each site (Fig. 2A and B), though the genome-wide contribution of neighbor genotypes to herbivore damage was significant (Fig. 1B). This result indicated a polygenic basis for the neighbor effect on herbivore damage. Next, we examined whether associational resistance was more likely than associational susceptibility. We focused on the sign of the estimated neighbor genotype effects, 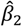, which represents positive or negative interactions between the two alleles of paired neighbors — i.e., associational resistance or susceptibility — against herbivore damage, respectively (*20, 23*). The top 0.1%-associated SNPs of the four phenotypes per site had both negative and positive 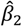 without clear bias (Fig. S6A and B). This result suggests that associational resistance and susceptibility are both possible, motivating us to examine the top-scoring SNPs with signs of neighbor genotypic effects *β*_2_ and other signatures.

**Figure 2.**
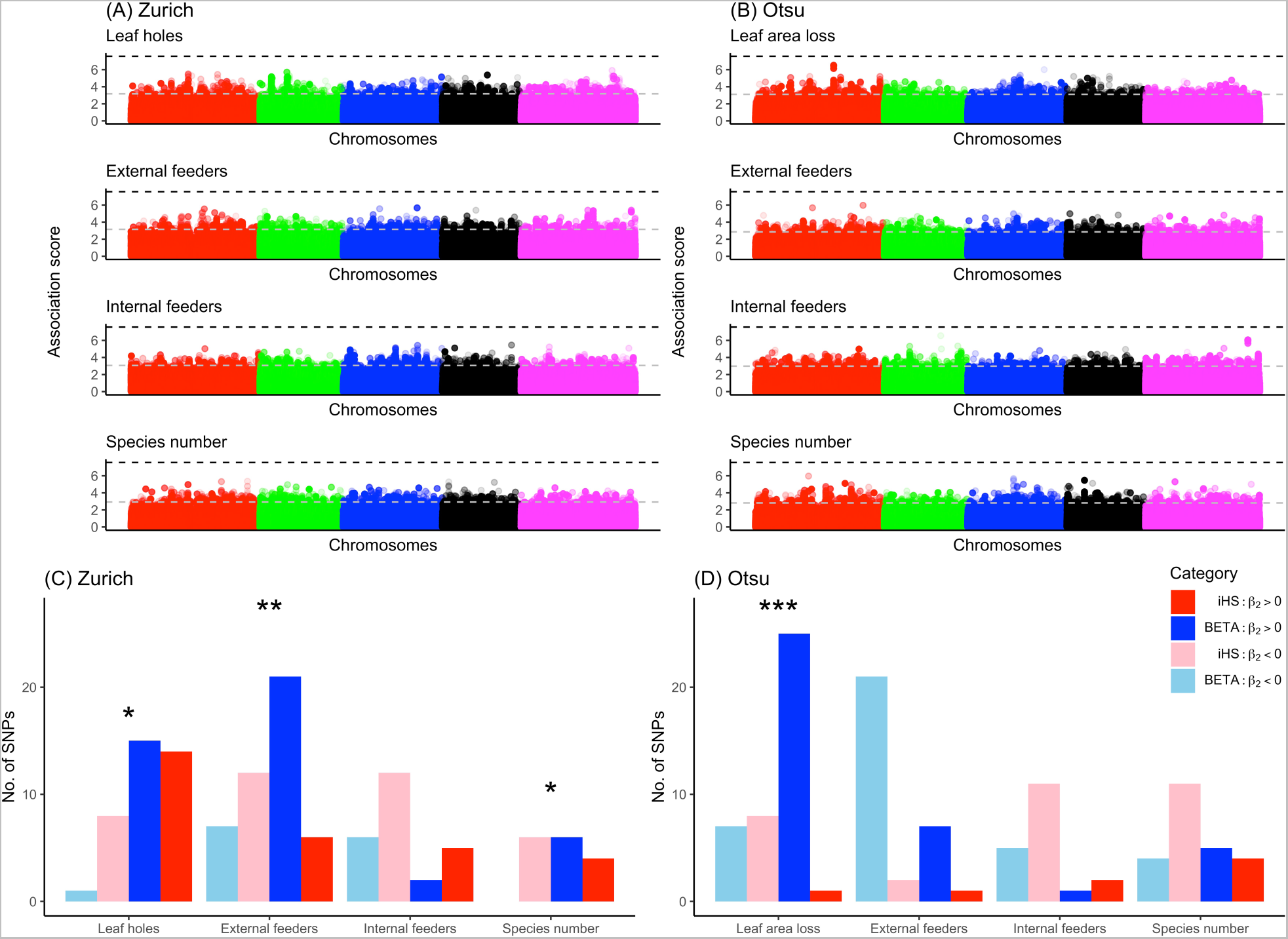
Genomic basis of neighbor effects on herbivore damage and community composition on *Arabidopsis thaliana* genotype mixtures. (A and B) Manhattan plots showing the −log_10_(*p*) association score of the neighbor genotype effect *β*_2_ across five chromosomes of *A. thaliana* at Zurich or Otsu. The horizontal dashed lines indicate the Bonferroni threshold at *p* = 0.05 (black) or the top 0.1% threshold of the association score (gray). (C and D) The number of SNPs shared between the selection scan (top >5%) and Neighbor GWAS (top >0.1%). The blue and red bars indicate balancing (BETA; blue) and directional selection (iHS; red) indices with positive (darker colors) or negative (paler colors) 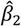, respectively. Asterisks indicate a significant excess of SNPs under balancing selection between positive and negative 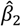; *** *p* < 0.001; ** *p* < 0.01; * *p* < 0.05 by Fisher tests.

To infer evolutionary patterns from the polygenic neighbor effects, we further analyzed the signature of natural selection on the top 0.1% SNPs relevant to associational resistance or susceptibility. Associational resistance and susceptibility represented by positive and negative 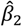 corresponds to negative and positive frequency-dependent selection on each SNP (*23*) (see also Supplementary Materials and Methods 2.1 and 2.4), and thereby are hypothesized to balance and unbalance multiple alleles at a locus, respectively (*12, 24*). We compared genome-wide signatures of balancing selection with those of directional selection to test whether balancing selection is more likely associated with positive 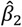. Herbivore damage at both sites and two further phenotypes in Zurich had more SNPs under balancing selection and associational resistance 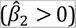 compared with those under directional selection and associational susceptibility 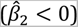 (one-sided Fisher tests, *p* < 0.05: Fig. 2C and D; Fig. S6). In contrast, none of the measured phenotypes showed opposite combinations i.e., a significant excess of SNPs under directional selection and associational resistance over those under balancing selection and associational susceptibility (one-sided Fisher test, *p* > 0.05). These patterns are consistent with the hypothesis that associational resistance can exert balancing selection on its responsible polymorphisms, highlighting the evolutionary background of polygenic neighbor effects.

The polygenic neighbor effects (Fig. 2A and B) made it difficult to identify important SNP predictors. We solved this problem using a genomic prediction approach (*25*) that incorporated all SNPs together for phenotype prediction. To predict the neighbor effects on herbivore damage of focal plants, we included all 1.2 million SNPs representing focal genotypes and neighbor genotypes in the least absolute shrinking and selection operator (LASSO) (*26*). With or without neighbor genotypes, LASSO prediction was validated using a test dataset collected in another year. Among the four phenotypes we had measured per site, the test dataset of herbivore damage in Zurich was slightly better explained by the neighbor-including LASSO than by the neighbor-excluding LASSO (Spearman’s *ρ* = 0.416 and 0.391, respectively: Fig. S7). This result indicates that herbivore damage can be better predicted by incorporating neighbor genotypes.

Using neighbor genotypes as a better predictor of herbivore damage in Zurich (Fig. S7A), we attempted to predict associational resistance or susceptibility to herbivore damage by specialist flea beetles. We did this by extrapolating the neighbor-including LASSO model to monoculture or mixture conditions *in silico*. From the neighbor-including LASSO, we extracted 756 neighbor-related SNPs to extrapolate the herbivore damage in Zurich (Fig. S8A and B). Assuming virtual mixture (a pair of two different genotypes) or monoculture (a pair of the same genotypes) conditions, we estimated the effects of two-genotype mixtures on the herbivore damage (Fig. S8C). This pairwise effect size had a negative mode in its distribution (Fig. 3A), suggesting the prevalence of associational susceptibility among the 199 genotypes. Furthermore, we found a significant negative correlation between this pairwise effect size and estimated herbivore damage under monoculture (*r* = *–*0.37; *p* < 0.001: Fig. S8F), indicating that susceptible plant genotypes impose more damage on their counterparts when planted with another genotype. Based on the pairwise effect size of the mixed planting (Fig. 3A), our simulations also confirmed that herbivore damage increased with a random increment in plant genotypic diversity (Fig. 3B; Fig. S8G). These results agree with those of a previous meta-analysis that reported negative effects of plant genotypic diversity on resistance to specialist herbivores (*9*). In this situation, we asked whether it would nevertheless be feasible to identify genotype pairs that would result in associational resistance at the stand level.

**Figure 3.**
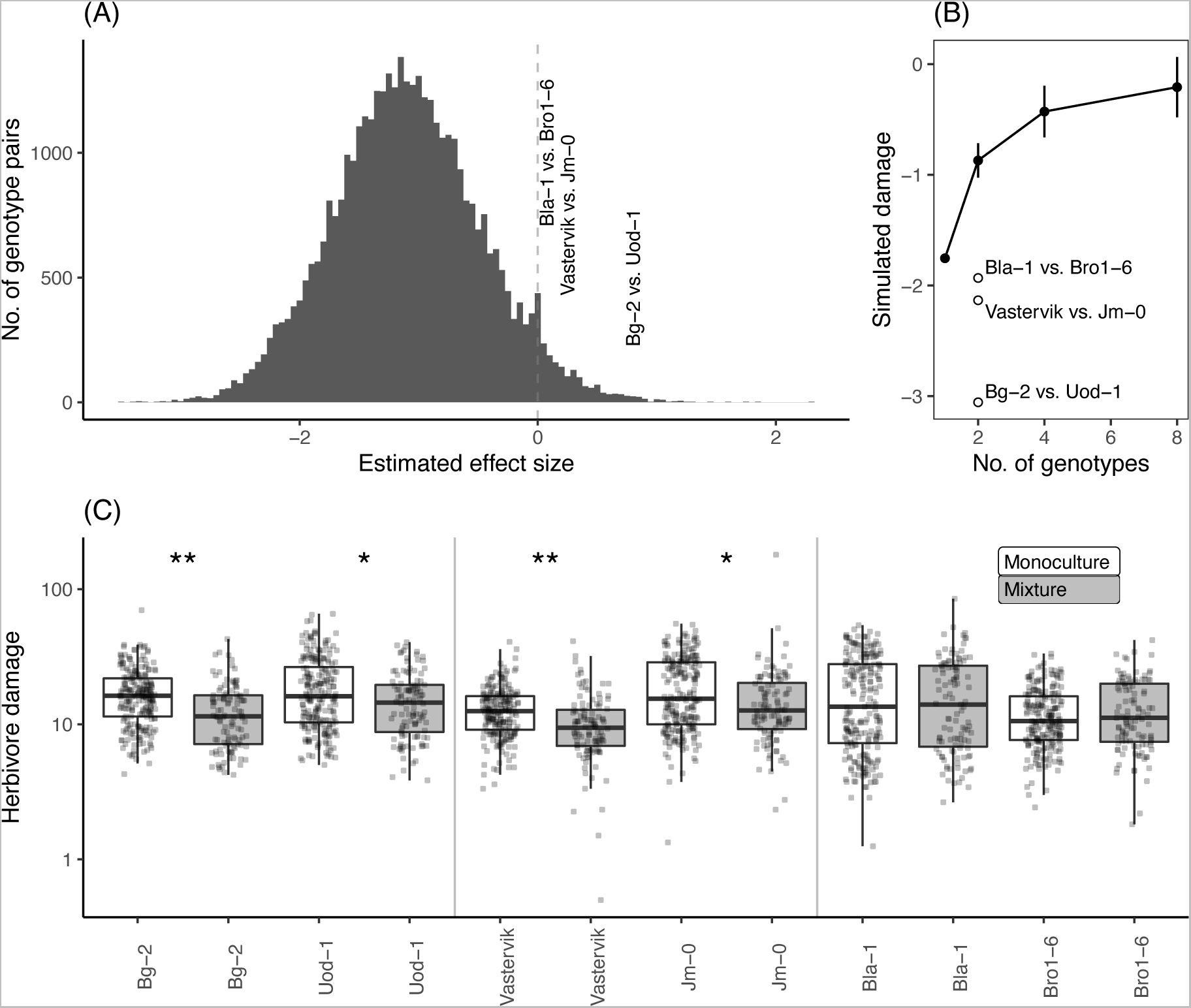
Effects of mixed planting on herbivore damage in silico and in situ. (A) Effect size estimates for pairwise mixed planting among the 199 *Arabidopsis thaliana* genotypes. Positive and negative values indicate associational resistance and susceptibility to herbivore damage, respectively. (B) Simulated damage (mean ± SD) is plotted against the number of randomly selected genotypes. (C) Herbivore damage by flea beetles on the three pairs of genotypes under monoculture (white) or mixture (gray) conditions in the Zurich field site. The y-axis represents the number of leaf holes divided by initial plant size (no./cm). Asterisks indicate significant differences in marginal means between the monoculture and mixture conditions (Table S5B): * *p* < 0.05 and ** *p* < 0.01.

Despite the prevalence of negatively interacting pairs (<0 in Fig. 3A), 823 pairs had a positive estimate of mixed planting (>0 in Fig. 3A). To verify associational resistance at the stand level *in situ*, we planted three genotype pairs under monoculture and mixture conditions at the Zurich site (Fig. S9). From the range of positive effect sizes (>0 in Fig. 3A), we focused on Bg-2 and Uod-1 as a pair with a large positive effect (effect size = 0.8); Vastervik and Jm-0 as a pair with a moderate positive effect (0.23); and Bro1-6 and Bla-1 as a pair with a slight positive effect (0.1). Consistent with this order of effect size, Bg-2 and Uod-1 indeed showed a significant reduction in herbivore damage in the mixtures in the field (Fig. 3C; Table S5; Table S6). Vastervik and Jm-0 also showed a significant reduction in herbivore damage in the mixture compared with the average monocultures (Fig. 3C; Table S5; Table S6). Expected from their smallest effect size, Bla-1 and Bro1-6 did not show a significant reduction in herbivore damage in the mixtures (Fig. 3C; Table S5; Table S6). In addition to field evidence, we allowed black flea beetles to feed on the three pairs in the laboratory. This additional experiment found significant differences in herbivore damage between Bg-2 and Uod-1 (likelihood ratio test, *p* < 0.01); and between Vastervik and Jm-0 (*p* < 0.05); but not between Bla-1 and Bro1-6 (*p* = 0.35; Fig. S10; Table S7), indicating that the least successful pair in the field could not alter herbivore damage even in a small-scale experiment. Field experiments and additional laboratory evidence have demonstrated that candidate genotype pairs underpin associational resistance to herbivory.

To understand the potential mechanisms of mixed planting, we also performed gene ontology enrichment analyses for the LASSO-selected SNPs relevant to associational resistance (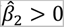; Table S8). We detected a significant enrichment of genes related to the jasmonic acid biosynthetic process (false discovery rate < 0.05; Table S9A), including the *LIPOXIGENASE2* (*LOX2*) and *LOX6* genes (Table S8). In contrast, jasmonate-related annotations did not appear when gene ontology analysis was applied for LASSO-selected SNPs relevant to associational susceptibility (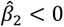; Table S9B). These results suggest that jasmonate-mediated defense signaling may partly explain associational resistance to flea beetles. *LOX2* is particularly known as an essential gene for the production of green leaf volatiles (*27*), which can reduce herbivory on neighboring plants (*15*). While the complex polygenic basis of neighbor effects makes it difficult to identify large-effect genes, comprehensive mutant analyses are needed to isolate causative genes.

Our study provides a proof-of-concept to predict impacts of intraspecific mixed planting on ecologically important phenotypes. Given that associational resistance has been widely reported in grasslands and forests (*11, 13*), the present findings highlight the potential ecological and evolutionary mechanisms of the effects of genetic diversity on plant resistance in terrestrial ecosystems. In addition to ecological interests, intraspecific mixed planting is also of applied interest because it may enhance plant resistance without complicating agronomic management (*17, 28, 29*). The genotypes of our key pair Bg-2 / Uod-1 are known to have similar flowering time (46.2 days for Bg-2 and 45.6 days for Uod-1 under a long-day condition) (*30*). This fact indicates that intraspecific mixed planting can be achieved without differentiating plant life cycles that may affect the timing of harvest (*28*). This novel strategy to identify genotype pairs with beneficial mixture effects may be more widely applicable to genotype mixtures in crops and other plantations.

## Supplementary Materials and Methods

### 1. Field GWAS experiments

#### 1.1. Plant genotypes

We used *Arabidopsis thaliana* genotypes that were selfed and maintained as inbred lines, called “accessions.” To study the genomic variation responsible for biotic interactions, we overlapped our accessions with those used in GWAS of microbial communities (*31*) and glucosinolates (*32*). We used 199 accessions with a few additional accessions (Table S1), all of which were genotyped by the RegMap (*18*) and 1001 Genomes (*19*) projects. Seeds of these accessions were obtained from the Arabidopsis Biological Resource Center (https://abrc.osu.edu/). The Santa-Clara accession was replaced with Fja1-1 in 2018 because the genotype of Santa-Clara was unavailable. For the genotype data, we downloaded a full imputed SNP matrix of 2029 accessions from the AraGWAS Catalog (*33*). Of the full 10,709,466 SNPs, we used 1,819,577 SNPs with minor allele frequency (MAF) at > 0.05. Our previous study detected the single-gene effects of *GLABRA1* (*GL1*) on flea beetle resistance (*21*); thus, L*er*(*gl1-1*) and Col(*gl1-2*) were included to test whether our GWAS experiments worked well. The L*er* or Col genome was assigned to the two *gl1* mutants, with only the *GL1* locus differing between the parental wild-type and *gl1* mutants.

#### 1.2. Field setting

To investigate two distinct herbivore communities, we used field sites within or outside a natural distribution range of *A. thaliana*. As a native site, we used the outdoor gardens of the University of Zurich-Irchel campus (Zurich, Switzerland: 47° 23’N, 8° 33’E, alt. ca. 500 m) (Fig. 1A). As an exotic site, we used the Center for Ecological Research, Kyoto University (Otsu, Japan: 35° 06’N, 134° 56’E, alt. ca. 200 m) (Fig. 1A). In the Otsu site, weeds were mown before the experiment, and the surroundings were covered with agricultural sheets before the experiment (Fig. 1A). In the Zurich site, each experimental block was placed in a separate bed (Fig. 1A) that was not accessible to molluscan herbivores.

Field experiments were conducted three times in 2017, 2018, and 2019. The field experiment at Otsu was conducted from late May to mid-June, and that at Otsu was conducted from late June to mid-July. The exact date of the field survey was annotated on the original data file (*34*). Plants were initially grown under controlled conditions and then planted in a field garden for three weeks. Seeds were sown on Jiffy-seven pots (33-mm diameter), and stratified under 4 °C for a week. Seedlings were cultivated for 1.5 months under a short-day condition (8 h light: 16 h dark, 20 °C). Plants were then separately potted in plastic pots (6 cm in diameter) filled with mixed soil of agricultural composts (Profi Substrat Classic CL ED73, Einheitserde Co. in Zurich; Metro-mix 350, SunGro Co., USA in Otsu) and perlites at a 3:1 L ratio. Covered with agricultural shading nets, the potted plants were acclimated to field conditions for a few days. A set of the 199 accessions and an additional Col-0 accession — namely, 200 individuals in total — was randomly assigned to each block without replacement and positioned in a checkered manner (Fig. 1C). Eight blocks of the 200 accessions were set at each site on 2017 and 2018 for GWAS, while the three replicates were set on 2019 for the model validation of LASSO (see “Modified Neighbor GWAS for LASSO” below). The blocks were >2.0 m apart.

#### 1.3. Phenotype survey

Insects and herbivorous collembola on individual plants were visually counted every 2– 3 days. These species were identified using a magnifying glass. Dwelling traces and mummified aphids were also counted as proxies for the number of leaf miners and parasitoid wasps, respectively. Eggs, larvae, and adults were counted for all species, as long as they could be observed by the naked eye. All counts were performed by a single observer (Y. Sato) during the daytime at each site. Small holes made by flea beetles were counted at the Zurich site and their maximum number throughout the experiment was used as an indicator of herbivore damage. This phenotyping was not applicable to Otsu, because the most abundant herbivores were not flea beetles. Instead, the percentage of leaf area loss was scored in Otsu at the end of the experiment as follows: 0 for no visible damage, 1 for <10%, 2 for >10% and <25%, 3 for >25% and <50%, 4 for >50% and <75%, and 5 for >75% of area eaten.

We also recorded the initial plant size and the presence/absence of inflorescences to incorporate these phenotypes as covariates in the statistical analyses. Initial plant size was evaluated by the length of the largest rosette leaf (mm) at the beginning of the field experiment because this parameter represents the plant size at the growth stage. The presence/absence of inflorescences was recorded 2 weeks after transplantation. Herbivore damage was evaluated by the number of leaf holes in Zurich, and the leaf area loss in Otsu as described above. The maximum number of individuals in each experiment was used as an index of the abundance of each insect species.

In this study, we defined indices of community composition based on herbivore feeding habits and species richness. Ordination analysis using the rda function of R (*35*) showed that community composition more significantly differed between the two sites than between 2017 and 2018 (redundancy analysis, *F* = 401, *p* < 0.001 for the sites; *F* = 152, *p* < 0.001 for the years: Fig. S1A); thus, we separated the dataset into Zurich and Otsu. The number of external or internal feeders was defined as the total number of individuals of leaf-chewing species (e.g., beetles and caterpillars) or species eating internal parts of a plant (e.g., phloem-sucking aphids, cell content-sucking thrips, and leaf miners). Because generalist herbivores were much fewer than specialist herbivores at both the sites (Fig. S1; Table S2), specialist-generalist classification was not applicable to our dataset. Carnivorous insects (e.g., parasitoid wasps and aphidophagous ladybirds) were also found but were much fewer than herbivores. The herbivore-carnivore ratio was thus not applicable, although these carnivorous insects were taken into consideration for insect species diversity. For the index of insect species diversity, we calculated the exponential Shannon diversity and Simpson diversity indices in addition to the total number of species i.e., species richness. However, Shannon diversity and Simpson diversity showed a discrete distribution that did not suit GWAS, and only the total number of species had quantitative phenotype values (Fig. S2). We therefore used the total number of species as an index of insect species diversity. The analysis of insect communities was performed using the vegan package (*35*) in R. All phenotypes except for the leaf area loss were ln(*x*+1)-transformed to improve normality for GWAS and genomic prediction. Unless otherwise stated, all figure presentations and basic statistical analyses were performed using R version 3.6.1 (*36*).

### 2. GWAS with focal and neighbor genotypic effects

#### 2.1. Neighbor GWAS model

To incorporate neighbor genotype identity into GWAS, we used a linear mixed model that included an additional fixed and random effect, called Neighbor GWAS (*20*). The core idea of this Neighbor GWAS method was inspired by the Ising model of ferromagnetism to estimate its interaction coefficient based on the genetic similarity between neighboring individuals (*20*). Let *x_i_* denote the allelic status at each SNP of the *i*-th focal plant and the *j*-th neighboring plants. The inbred accessions took two states as *x_i_ ∈* {-1, +1}. A phenotype value of the *i*-th focal individual plant *y_i_* was then given as

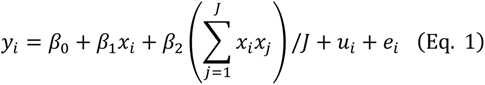

where *β*_0_ is the intercept; *β*_1_*x_i_* is a fixed effect of the focal genotype and the same as standard GWAS; and the second coefficient *β*_2_ determines positive or negative effects from the mean allelic similarity 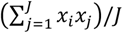 at a given locus between the focal individual *i* and neighboring individuals *j* up to the total number of neighboring individuals *J*. The random effects *u_i_* and residuals *e_i_* follow a normal distribution as 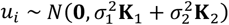 and 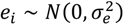, where 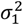 and 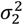 indicated the variance component parameters for the polygenic effects from focal and neighbor genotypes, respectively. **K**_1_ or **K**_2_ represents a kinship matrix among *n* plants given by the cross-product 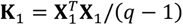 or 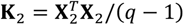, where **X**_1_ or **X**_2_ was a *q* × *n* matrix that includes all focal genotype values or neighbor genotype similarity, respectively. A standard GWAS model is a subset of the Neighbor GWAS model (Eq. 1). When *β*_2_ and 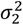 was set at 0, the Neighbor GWAS model was equivalent to the standard GWAS model. In the context of the magnetic model, positive or negative *β*_2_ determines whether neighbor clustering or mixture can maximize phenotype values at the population level, respectively (*20*). In the context of the population genetic model, the positive or negative *β*_2_ respectively represent symmetric positive or negative frequency-dependent selection that increases or decreases mean fitness at an intermediate frequency of the two alleles, respectively (*23, 37*). In the case of plant defense, herbivory corresponds to negative effects on plant fitness. In contrast to the interpretation of frequency-dependent selection on fitness, positive *β*_2_ represents a positive interaction that decreases the negative effects on fitness, whereas negative *β*_2_ represents a negative interaction that increases the negative effects on fitness. In our study, SNPs with positive *β*_2_ had the potential to drive positive interactions that could reduce herbivore damage by mixing two alleles.

#### 2.2. PVE and association mapping

Using the Neighbor GWAS model (Eq. 1), we estimated the proportion of phenotypic variation explained (PVE) by genetic factors and performed association mapping of the SNP marker effects. The statistical significance of the variance components, 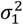 and 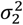, or the fixed effects, *β*_1_ and *β*_2_, was determined by likelihood ratio tests between models with or without a single parameter. The proportion of phenotypic variation explained (PVE) by the two genetic factors was defined as 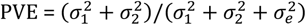. The genomic heritability in the standard GWAS was given by 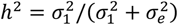 when 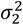 was set to 0. Linear mixed models with variance component parameters 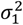 and 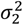 were solved using the average information-restricted maximum likelihood method (*38*). To perform association mapping, we then tested single-marker effects *β*_1_ and *β*_2_ using eigenvalue decomposition on a weighted kinship matrix 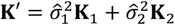 (*38*). The likelihood ratio was used to calculate *p*-values of each parameter based on *χ*^2^ distribution with one degree of freedom. This line of GWAS analysis was implemented in the rNeighborGWAS (*20*) package, which internally uses the gaston package (*38*).

To determine the space of neighbor effects, we conducted variation partitioning and association mapping at *J* = 4 (up to the nearest neighbors) and *J* = 12 (up to the second nearest neighbors). Starting from the smallest space, our previous simulations showed that the optimal balance between false positive and negative detection of causative SNPs was achieved when phenotypic variation explained by neighbor effects turned significant (*20*). To anticipate this notion, we broadened the reference space of the neighbor effects to the second-nearest neighbors i.e., *J* = 12. This association mapping at *J* = 12 found significant SNPs regarding the leaf holes and leaf area loss (Fig. S4C and Fig. S5C); however, the positions of the peaks were different from those of *J* = 4. Furthermore, the neighbor effects on these phenotypes at *J* = 12 exhibited inflated *p*-values (see quantile-quantile plots in Fig. S4C and Fig. S5C), indicating the risk of false positives. The line of results at *J* = 12 indicates that the genomic basis of neighbor effects cannot be further resolved by incorporating long-range neighbor effects. We therefore presented the results of *J* = 4 in the main text, while including the results of *J* = 12 for phenotypic variation (Fig. S3), association mapping (Fig. S4 and S5), and selection scans (Fig. S6) in the Supplementary Figures and Tables.

#### 2.3. Post-GWAS analysis (i): List of candidate genes

Candidate genes near SNPs with the top 0.1% *p*-values were searched within 10 kbp around each SNP after association mapping. Functional annotation data from The Arabidopsis Information Resource (TAIR) were used for the gene model and description of *A. thaliana* (*39*).

#### 2.4. Post-GWAS analysis (ii): Selection scan

To test whether associational resistance and susceptibility coincided with the signatures of selection, we used two methods that detect balancing or directional selection based on a sweep pattern near the target SNP (*40, 41*). First, the signature of directional selection was analyzed using extended haplotype homozygosity (EHH) and its integrated haplotype score (iHS), which were designed to detect positive selection for new mutations (*40*). We focused on such a positive selection for directional selective pressure because purifying selection i.e., negative selection results in monomorphism and thus is not applicable for polymorphic sites. The EHH and iHS were calculated using the rehh package (*42*). Second, the signature of balancing selection was analyzed using the BetaScan method, which detects allele frequency correlations near the target SNP (*41*). Default settings were applied to the rehh package and BetaScan methods. SNPs in the top 5% of the empirical distributions were considered to be those under selection. Ancestral alleles were determined in comparison with the whole genome sequence of *A. lyrata*. The multiple alignment FASTA file comparing *A. thaliana* and *A. lyrata* genome sequences was downloaded from the Ensembl database (ftp://ftp.ensemblgenomes.org/pub/plants). Fisher’s exact probability tests were applied for a 2 × 2 matrix that included the number of SNPs for balancing or directional selection; and for associational resistance 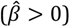 or susceptibility 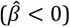 (Fig. 2C and D; Fig. S6E and F). One-sided Fisher tests were used to test the excess of balanced or positively selected SNPs. We also changed the threshold of the top-scoring SNPs for Neighbor GWAS at 0.5% and 1%, but these thresholds did not alter our conclusion in the main text (results not shown).

### 3. LASSO with focal and neighbor genotypic effects

#### 3.1. Modified Neighbor GWAS for LASSO

To perform multiple regressions on all SNPs, we used sparse regression that could simultaneously select important SNP predictors and estimate their coefficients. The Neighbor GWAS model (Eq. 1) is expressed as a multiple regression model, as follows:

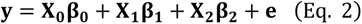

where *y* is a phenotype vector; **β_0_** is a vector including coefficients for an intercept and non-genetic covariates; **β_1_** and **β_2_** are vectors including coefficients of focal and neighbor genotype effects, respectively; **X**_0_ is a matrix that includes a unit vector and non-genetic covariates for *n* individuals. **X**_1_ is a matrix that includes the focal genotype values for *n* individuals and *q* SNP markers. **X**_2_ is a matrix that includes the neighbor genotype similarity for *n* individuals and *q* SNP markers as follows:

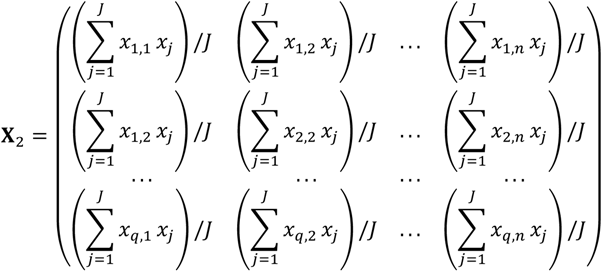

To simultaneously perform variable selection and coefficient estimation, we applied the least absolute shrinkage and selection operator (LASSO) (*26*) to Eq. 2. Because LASSO is sensitive to high correlations among explanatory variables, we further cut off 1,819,577 SNPs to 1,242,128 SNPs with the criterion of linkage disequilibrium (LD) at *r*^2^ < 0.8 between adjacent SNPs. The initial plant size, presence/absence of inflorescences, and experimental blocks were considered as fixed covariates. Important variables were selected from 1,242,128 SNP markers and the same number of neighbor-related SNPs using LASSO. We used the Python version of the glmnet package (*43*) to perform LASSO. The kinship or population structure among individuals was implicitly considered because LASSO regression could deal with all the SNPs simultaneously. While a gradient of sparse regressions from the LASSO, via the elastic net, to the ridge regression was available in the glmnet package (*43*), we used the sparsest regression, LASSO, because of a computational burden of recursive calculation during the effect size estimation and simulation (see “Effect size of mixed planting” below).

To determine the LASSO regularization parameter *λ*, we first trained the LASSO models with the learning data (years 2017 and 2018) and then validated their outputs using the test dataset collected in another year (i.e., 2019; see also “Field setting” above). The predictability of the four phenotypes was evaluated based on the correlations between the predicted and observed values of each phenotype. Spearman’s rank correlation *ρ* was used because some phenotypic values were not normally distributed. The predicted values were obtained from the LASSO models with different values of *λ*. To assess genetically based predictability, we quantified observed phenotype values in 2019 as residuals of a standard linear model. This standard linear model incorporated the same non-genetic explanatory variables as the LASSO model, including the initial plant size, presence of inflorescence, and difference in three experimental blocks, while each phenotype was considered a response variable. To determine whether the incorporation of neighbor genotypes improved the correlation with the test data, we compared LASSO with or without neighbor genotypes across a series of *λ*. If the neighbor-including LASSO yielded a larger correlation than the neighbor-excluding LASSO at a given *λ*, this indicates that neighbor genotypes were able to improve the predictability of a target phenotype by LASSO. In this context, the maximum *ρ* of the neighbor-including LASSO was larger than that of the neighbor-excluding LASSO on herbivore damage in Zurich (Fig. S7). Furthermore, the neighbor-including LASSO achieved this maximum *ρ* even at stringent regularization (= larger *λ*) compared to the neighbor-excluding LASSO (Fig. S7A). For the Otsu site, the neighbor-including LASSO also had slightly larger correlations with herbivore damage than the neighbor-excluding LASSO, supporting the improved predictability of herbivore damage by neighbor genotypes at another site (Fig. S7B). None of the community composition phenotypes, however, showed better predictability by the neighbor-including LASSO (Fig. S7B). This was presumably because the abundance of the predominant species differed between study years (Fig. S1B-G). These additional results support the improved predictability of herbivore damage but suggest difficulty in predicting community composition by neighbor genotypes.

When the neighbor-including LASSO outperformed the neighbor-excluding ones at a given *λ*, we obtained the vectors of the estimated coefficients 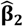 that were able to improve the phenotype prediction. LASSO could yield multiple sets of 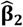 across a series of *λ* where the neighbor-including LASSO yielded larger correlations. Larger *λ* tend to give fewer non-zero SNPs with large coefficients, while smaller *λ* tend to give more non-zero SNPs with small coefficients. To consider the polygenic basis of neighbor effects, we averaged the estimated coefficients 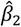 per SNP across the range of *λ*, resulting in 756 SNPs with non-zero *β*_2_ for the herbivore damage in Zurich (see the main text). This estimated vector of neighbor coefficients 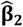 was used to estimate the effect size.

#### 3.2. Post-LASSO analysis (i): The effect size of mixed planting

To estimate the pairwise effect size of mixed planting, we extrapolated the LASSO models Eq. 2 under a virtual monoculture (= a pair of the same accession) or pairwise mixture (= a pair of different accessions). The pairwise effect size was determined by the difference in the linear sum 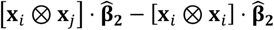 between a pair of accessions. The first term 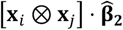 represents the phenotype values expected from different genotype vectors between the accession *i* and *j* (= pairwise mixture), whereas the second term 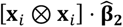 represents those expected from the same genotype vectors between the accession *i* and *i* (= monoculture). The element-wise product [**x***_i_* ⊗ **x***_j_*] or [**x***_i_* ⊗ **x***_i_*] represents the neighbor genotype similarity between a pair of different or the same accessions, respectively. Because the neighbor genotype effects turned out to have a polygenic basis (Fig. 2A and B), the genotype pairs predicted by many moderate-effect loci were suitable for testing the estimated effects of mixed planting. In contrast, genotype pairs showing the largest effect size were selected based on a few large-effect but less reliable loci. Assuming that multiple moderate-effect loci could result in the effects of mixed planting, we avoided the extreme tail of the effect size distribution when focusing on pairs. Also note that *β*_2_ in the neighbor GWAS models (Eqs. 1 and 2) denotes symmetric interactions between the focal *i* and neighbor *j* individuals (*20*), and thereby [**x***_i_* ⊗ **x***_j_*] and [**x***_j_* ⊗ **x***_i_*] have the same effects on a target phenotype. Even when asymmetric effects are incorporated, they do not affect the *relative* differences in phenotype values between *i* and *j* (*23*). Thus, we focused on the symmetric neighbor effects *β*_2_ to estimate the relative effect size of a pairwise mixture on a phenotype *y*.

To test whether the increasing number of plant genotypes increases or decreases herbivore damage, we also simulated herbivore damage in Zurich — i.e., ln(no. of leaf holes+1) — using the estimated vector of the neighbor coefficients 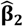. Assuming the nearest neighbors in a two-dimensional lattice, we simulated mixtures of up to eight genotypes. The herbivore damage was predicated by its marginal value with respect to the net neighbor effects 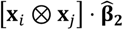. To examine the overall and selected patterns, we tested two types of genotype selection: (i) random selection from all pairs or (ii) random selection from pairs with positive estimates of pairwise mixed planting (positive values in Fig. 3A). First, eight genotypes were randomly selected out of the 199 accessions to represent overall pattern (Fig. 3B). We listed one (monoculture), two, four, or eight (full mixture) genotype combinations among the selected eight genotypes, and averaged their predicted damage 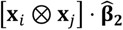 among all the combinations. Second, four positively interacting pairs (Fig. 3A) were randomly selected to test whether random selection of positive pairwise interactions could yield positive relationships between genotype number and anti-herbivore resistance (Fig. S8D). Duplicates of accessions were not allowed when selecting the four pairs of two paired accessions. This line of random sampling was performed 9999 times to calculate the mean and standard deviation. In the first case, Figure 3A shows a negative relationship between the number of genotypes and plant resistance. In the second case, herbivore damage decreased by paired mixing but increased by four- and eight-genotype mixing (Fig. S8D). This was because scaling up pairwise mixtures to four or eight genotypes confounded negatively interacting pairs. In addition to Figure 3A, these supplementary results also support the difficulty in targeting positive relationships between genotype richness and anti-herbivore resistance.

To determine whether geographical or genomic similarity could also predict the pairwise effect size between *A. thaliana* accessions, we analyzed the correlations of the pairwise effects with either geographical or genetic similarity. The statistical significance of the Pearson correlation *r* between a pair of matrices was determined using Mantel tests implemented in the vegan package (*35*) with 999 permutations. The geographical distance was determined by the Euclidean distance between the latitude and longitude of the locality of each accession. The locality of *A. thaliana* accessions was obtained from the AraPheno database (*33*). The genetic distance between the two accessions was determined using a kinship matrix **K**_1_. This additional analysis confirmed that the pairwise effect sizes were not related to geographical or genetic distance (Mantel tests, *r* = 0.02, *p* = 0.309 for geographical distance; *r* = *–*0.007, *p* = 1 for genetic distance: Fig. S8E and F). These additional analyses indicate that the estimated effect size is predictable by neither genome-wide genetic similarity nor geographical origin between accessions.

#### 3.3. Post-LASSO analysis (ii): GO enrichment analysis

To infer a category of genes related to positive and negative neighbor effects, we performed gene ontology (GO) enrichment analyses for candidate genes near LASSO-selected SNPs (i.e., SNPs with non-zero 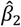). Same as the post-GWAS analysis above, we searched for genes within 10 kbp around each selected SNP. We then omitted duplicated genes after listing the candidate genes. We finally performed Fisher’s exact probability tests for each GO category against the entire gene set of *A. thaliana*. Multiple testing was corrected using the false discovery rate (FDR) (*44*). The entire set was built upon the TAIR GO slim annotation (*39*) using the GO.db package (*45*) in R. To summarize the results of the GO enrichment analysis, we applied the REVIGO algorithm (*46*) to the list of significant GO terms at FDR < 0.05. When summarizing the significant GO terms, we focused on the Biological Process with the similarity measure at 0.7 (i.e., the same as the default setting). We used the rrvgo (*47*) and org.At.tair.db (*48*) packages in R to run the REVIGO algorithm. This line of GO analysis was separately performed for SNPs that had negative or positive 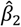 to detect GO terms unique to positive or negative neighbor effects on anti-herbivore resistance. Note also that post-GWAS GO analyses possess the issue of statistical non-independence due to linkage disequilibrium in standard GWAS (*49*). However, LASSO was unlikely to be subject to this issue because (i) this sparse regression could sparsely select SNP variables across a genome; (ii) we pruned adjacent SNPs on the strong LD at *r*^2^ > 0.8; and (iii) we focused on unique genes before using the Fisher tests. Therefore, we applied the conventional GO enrichment test based on the Fisher tests with FDR correction for the LASSO results. The in-house R package that includes utility functions of the GO enrichment analysis is available at GitHub and Zenodo (*50*).

### 4. Mixed planting experiment

#### 4.1. Field experiment

To test the effects of mixed planting on herbivore damage, we transplanted three pairs of accessions (i.e., Bg-2 and Uod-1; Vastervik and Jm-0; and Bla-1 and Bro1-6) under mixture and monoculture conditions. The theory of plant neighbor effects suggests that both plant patch size and neighbor composition should be manipulated to distinguish the effects of mixed planting from the density-dependent attraction of herbivores (*14*). We therefore set the large or small plant patches in addition to monoculture or mixture conditions. The field experiment was conducted from late June to July 2019 and 2021 in the outdoor garden of the University of Zurich-Irchel. Plants were first grown under short-day conditions and then transferred to an outdoor garden following the same procedure as the field GWAS above. Two accessions were then mixed in a checkered manner under the mixture condition, while either of the two accessions was placed under the monoculture conditions. The large patch included 64 potted plants in 8 × 8 trays and had a single replicate, while the small patch included 16 plants in 4 × 4 trays and had three replicates (photo of Fig. S9). In the mixture setting, the two potted accessions filled the square space in a checkered manner without a blank position (photographs in Fig. S9). The total number of initial plants was two accessions × three pairs × the mixture or monoculture × the large or small patches × two years = 2,016 individuals. Only a few pots per plot were labelled for tracking the plots in the field, whereas the other pots were not labelled to blind their information. The initial plant size was measured in the same manner as in the field GWAS experiment. Leaf holes were counted three weeks after transplantation. Four plants died during the field experiment, resulting in a final sample size of 2,012 plants.

#### 4.2. Statistical analysis

We used linear mixed models to analyze the number of leaf holes because this variable appeared to be normally distributed. The response variable was ln(x+1)-transformed number of leaf holes per plant to improve the normality. The explanatory variables were plant accession, mixture or monoculture condition, small or large patches, and study years. The initial plant size, represented by the length of the largest leaf (mm), was considered as an offset term. Two-way interactions were also considered among the plant accessions, mixture conditions, and patch conditions. Because the large and small patches had different numbers of individual plants, this imbalance was dealt with using a random factor. We split the large patch by 4 × 4 potted plants (= the same size as the small patch; see also a photo in Fig. S9), and considered these subplot differences — i.e., a total of 126 subplots — as a random effect. The significance of each explanatory variable was tested using Type III analysis of variance based on Satterthwaite’s effective degrees of freedom (*51*). To compare herbivore damage for each accession between the mixture and monoculture conditions, we calculated marginal means for the full model based on Satterthwaite’s method with Sidak correction for multiple testing (*52*). For these analyses of leaf holes, we used the lme4 (*53*), lmerTest (*51*), and emmeans (*52*) packages in R. Box plots visualize the median with upper and lower quartile, with whiskers extending to 1.5 × inter-quartile range.

To examine the effects of patch size and year in addition to mixed planting (Fig. 3C), we analyzed a separate dataset for patch conditions and study years (Fig. S9A-D; Table S6). Consistent with the order of the estimated effect size (Fig. 3A), the estimated marginal means across these conditions showed the largest sum of effects of mixed planting between Bg-2 and Uod-1 (= 0.495 in Table S5B) and the second largest effect between Vastervik and Jm-0 (= 0.453 in Table S5B). The significant effects of mixed planting on herbivore damage were more detectable in the large patches than in the small patches (Fig. S9). The Bg-2 or Uod-1 accessions showed a significant reduction in herbivore damage among five cases out of the two accessions × two years × two patch conditions (Fig. S9; Table S6) and a marginally significant case in the small patch (*p* = 0.053 in Table S6A). The Vastervik or Jm-0 showed three significantly positive cases favoring the reduction in herbivore damage out of the eight conditions (Fig. S9; Table S6), indicating less consistency than the Bg-2 and Uod-1 pairs under diverse conditions. The Bla-1 and Bro1-6 pairs did not have significantly positive cases favoring the reduction in herbivore damage out of the eight conditions and even had one case of increased damage by mixed planting (Fig. S9; Table S6). The main results and separate data show that the magnitude of the positive mixing effect is comparable to the order of the estimated effect size.

### 5. Laboratory choice experiment

#### 5.1. Insect materials

To examine feeding by flea beetles, we conducted laboratory choice experiments using one of the two major flea beetles, the black flea beetle *Phyllotreta astrachanica*. Adult *P. astrachanica* were collected from *Brassica* spp. at the University of Zurich-Irchel. Adults and larvae were reared on German turnips (Kohlrabi) following a previously established protocol (*54*). The species of flea beetles were identified by the DNA sequence of the mitochondrial gene encoding Cytochrome C Oxidase Subunit I (*COI*). DNA was extracted using ZYMO RESEARCH Quick-DNA Tissue/Insect Kits (cat. no. D6016). We used universal COI primers designed by Folmer et al. (*55*) for Polymerase Chain Reaction (PCR) amplification under the following conditions: Initial denaturation at 95 °C for 5 minutes followed by 40 cycles of 95 °C for 15 seconds, 50 °C for 30 seconds, 72 °C for 60 seconds and a final extension at 72 °C for 3 minutes. The PCR products were sequenced by Sanger sequencing. We compared our sequences with the COI sequences registered by Hendrich et al. (*56*), which included 15 *Phyllotreta* species with several individual vouchers per species collected in Central Europe. Our sequences and the registered sequences were clustered using a neighbor-joining tree and the default alignment method implemented in the Qiagen CLC Main Workbench. We identified species from our samples based on phylogenetic clusters. Our sequence data are registered in GenBank with IDs from OQ857829 to OQ857834, which include three individuals of black- and yellow-striped flea beetles.

#### 5.2. Experimental setting

We used three pairs of six *A. thaliana* accessions, Bg-2 vs. Uod-1, Vastervik vs. Jm-0, and Bla-1 vs. Bro1-6. Seeds were sown on Jiffy-seven pots (33-mm diameter) and stratified at 4 °C for a week. Seedlings were cultivated under long-day conditions (16 h light: 8 h dark, 22/20 °C) for 3 weeks, with liquid fertilizer added a week after the start of cultivation. We then allowed two adult beetles to feed on two individuals × two accessions for three days under long-day conditions. The feeding arena was constructed using a transparent plastic cup (129 mm in diameter and 60 mm in height) that enclosed four Jiffy-potted seedlings. Excluding cups without any infestation by *P. astrachanica*, we obtained 15-20 replicates of the feeding arena per pair.

#### 5.3. Statistical analysis

We analyzed the number of leaf holes per plant as a response variable. Because the number of leaf holes in this short-term laboratory experiment was zero truncated (Fig. S10), we used generalized linear models with a negative binomial error and log-link function. Plant accessions and arena IDs were included as the explanatory variables. Likelihood ratio tests based on a *χ*^2^-distribution were used after checking whether the ratio of residual deviance to the residual degree of freedom was nearly one. The significance of each explanatory variable was tested by excluding one variable from the full model. The glm.nb function in the MASS package in R was used for generalized linear models with negative binomial errors. Likelihood ratio tests showed that flea beetles showed a significant preference between Bg-2 and Uod-1; and between Vastervik and Jm-0; but not between Bla-1 and Bro1-6 (Table S7). The effect of the experimental area on leaf holes explained deviance but was only significant in the Bg-2 and Uod-1 pairs (Table S7).

## Acknowledgements

The authors thank K.K. Thomsen, L. Mohn, M. Brasser, and all members of Shimizu group for help with the field setup in Zurich; G. Yumoto, L.G. Kawaguchi for field assistance in Otsu; T. Tsuchimatsu for advice on GWAS during the early stage; M. Yamazaki for advice on the *Arabidopsis* cultivation and molecular experiments; F. Beran for advice on the barcoding of flea beetles; and J. Bascompte, M.A. Barbour, and S.E. Wuest for comments on the manuscript.

## Author contributions

Y.S.: conceptualization, project administration, investigation, data collection, formal analysis, funding acquisition, draft writing, reviewing and editing; R.S.I: investigation (field), reviewing and editing; K.T.: investigation (bioassay), reviewing and editing; B.S.: formal analysis (mixed planting), funding acquisition, reviewing and editing; A.J.N.: conceptualization, supervision, funding acquisition, reviewing and editing; K.K.S.: conceptualization, supervision, funding acquisition, reviewing and editing.

## Data availability

All the source codes and original data generated in this study are available in the GitHub repository (https://github.com/yassato/AraHerbNeighborGen). This repository is also deposited in Zenodo (https://doi.org/10.5281/zenodo.7945318) (*34*).

## Supplementary Figures and Tables

**Figure S1.**
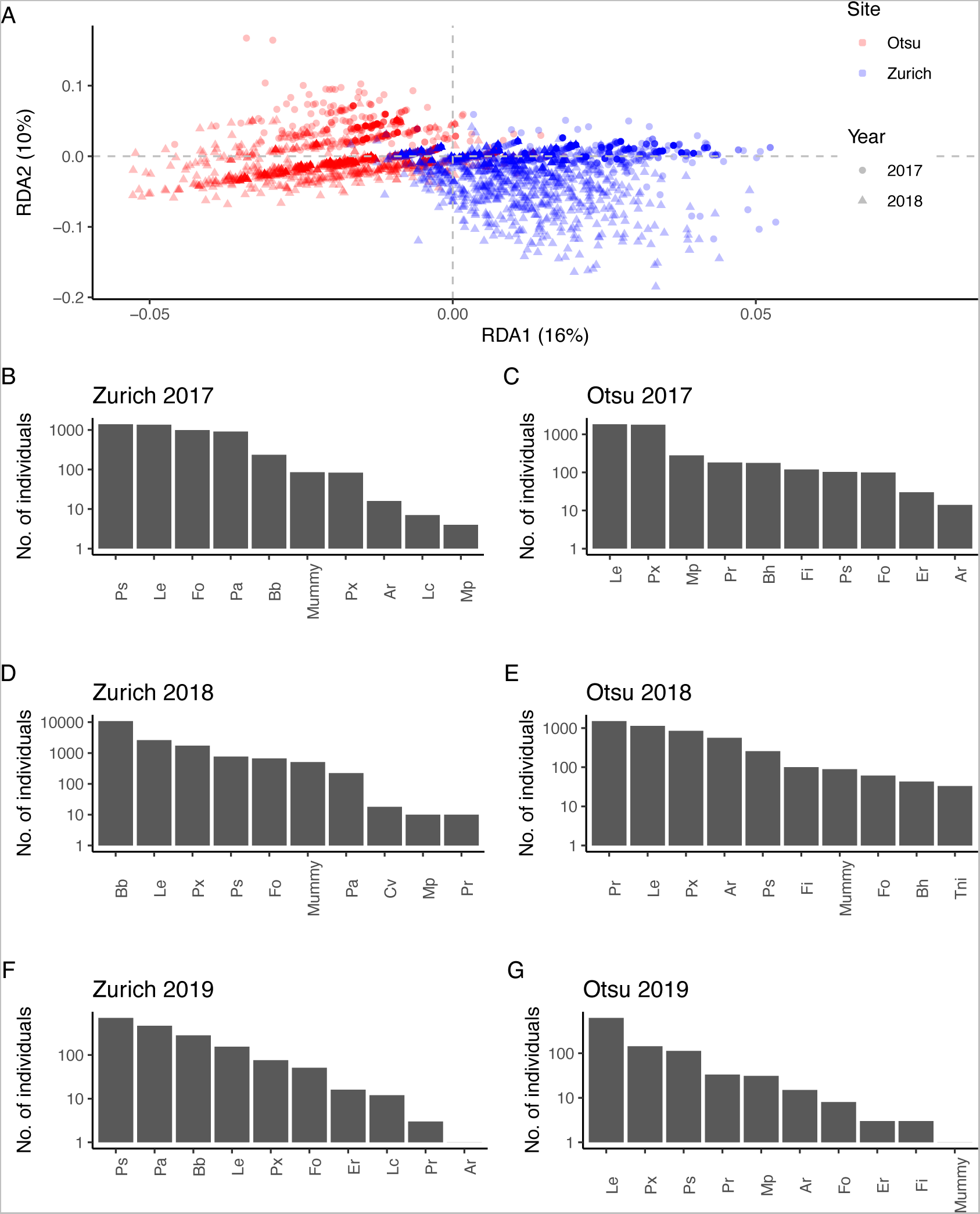
Insect communities observed on field-grown *Arabidopsis thaliana* in Zurich and Otsu. (A) Redundancy analysis showing the dissimilarity in insect communities between the two sites and years. The plot type indicates the study year and site: circles (2017), triangles (2018), blue (Zurich), and red (Otsu). The percentages of community variation explained by RDA1 and RDA2 are shown on each axis. (B-G) Abundance of major insect species observed from 2017 to 2019. The species name and its abbreviation correspond to those summarized in Table S2

**Figure S2.**
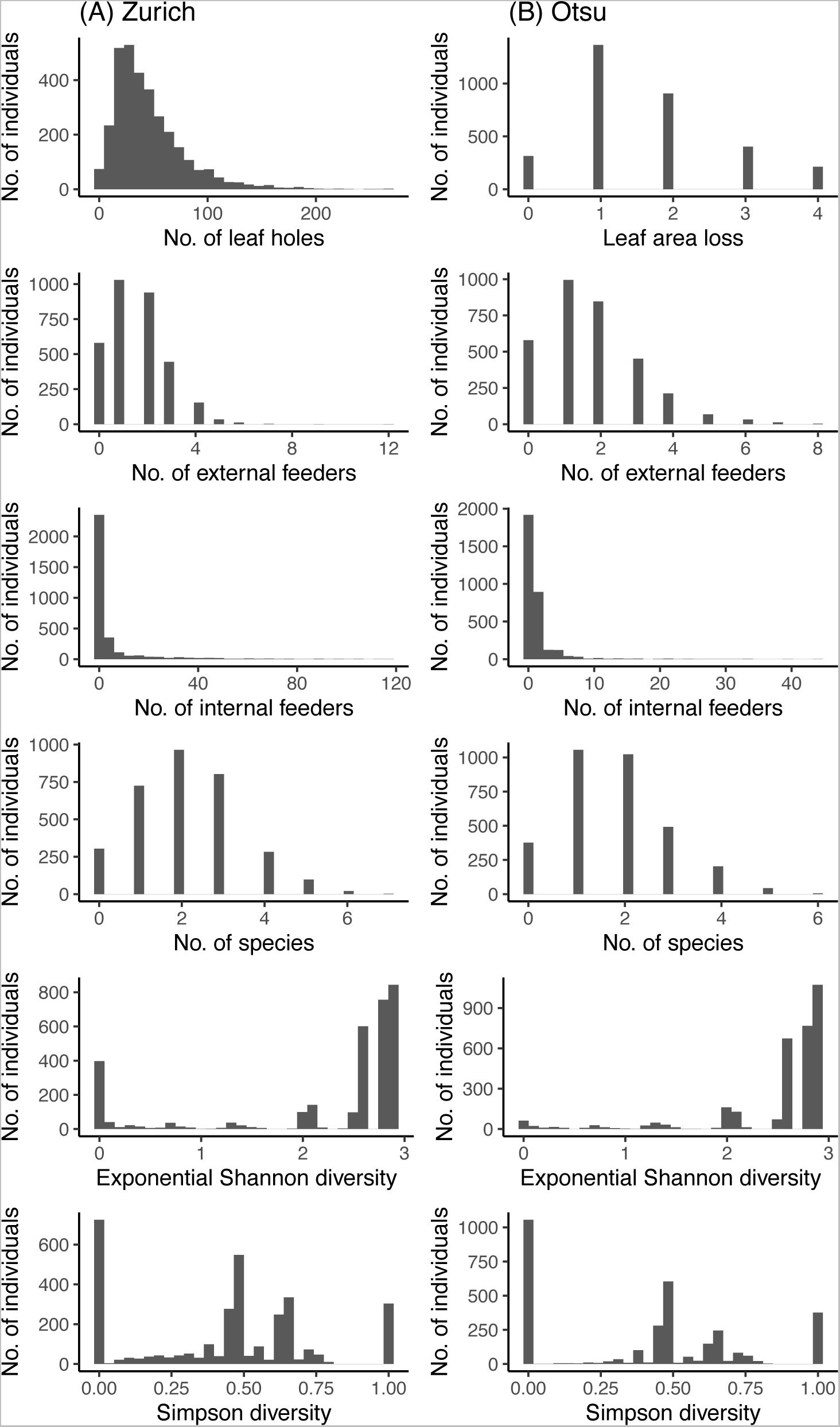
Histograms of phenotypic values per plant in Zurich (A) and Otsu (B). Shown are five phenotypes subject to GWAS (the upper four rows) as well as two measures of insect species diversity (the lower two rows).

**Figure S3.**
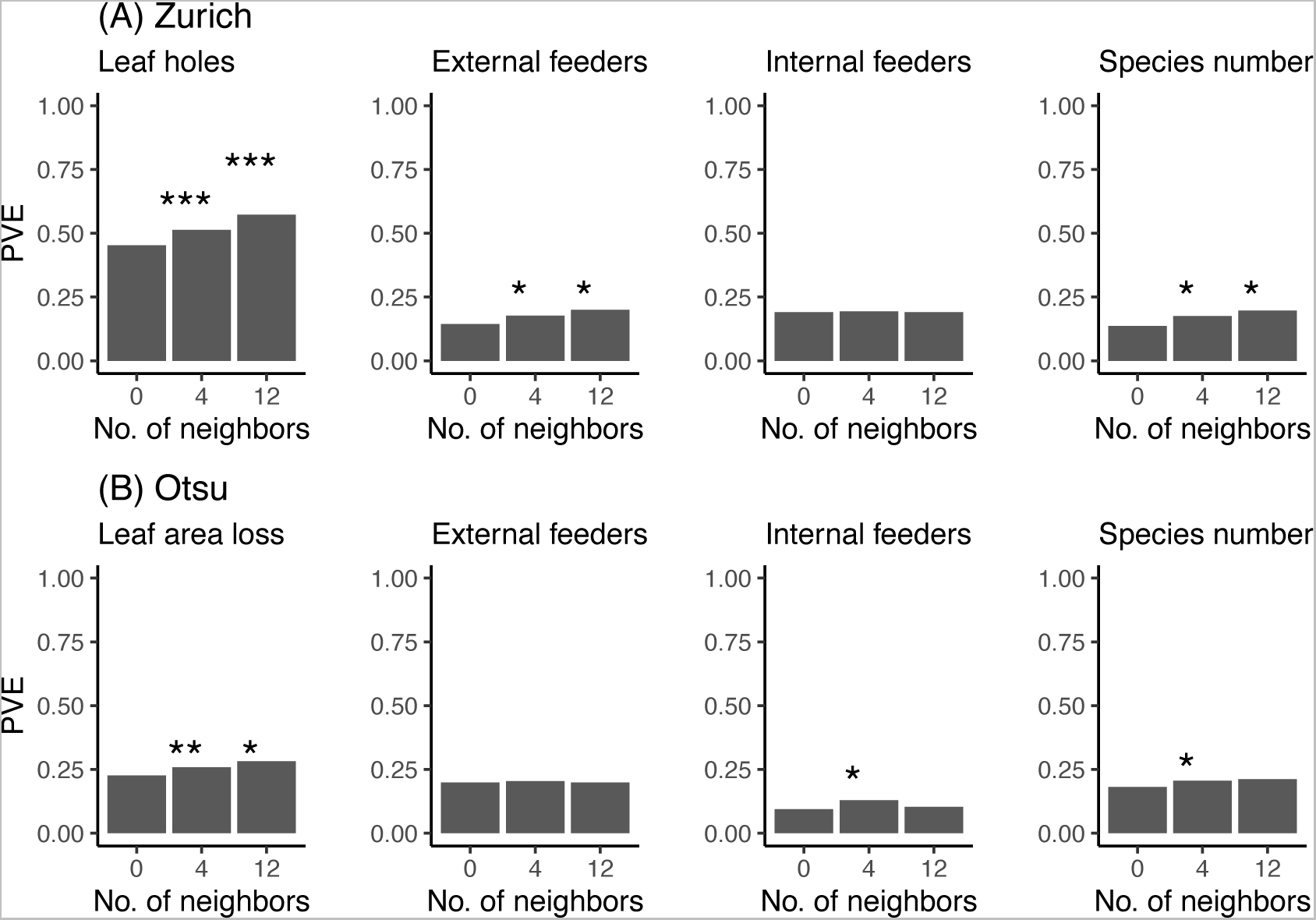
Proportion of phenotypic variation explained by the standard or Neighbor GWAS model in Zurich (A) or Otsu (B). The number of neighbors at 0 corresponded to the standard GWAS that quantified genomic heritability alone. The numbers of neighbors *J* at 4 and 12 correspond to the first and second nearest neighbors, respectively. Asterisks indicate the statistical significance of the neighbor GWAS (*J* = 4 or 12) over standard GWAS (*J* = 0): *** *p* < 0.001; ** *p* < 0.01; * *p* < 0.05 with likelihood ratio tests.

**Figure S4.**
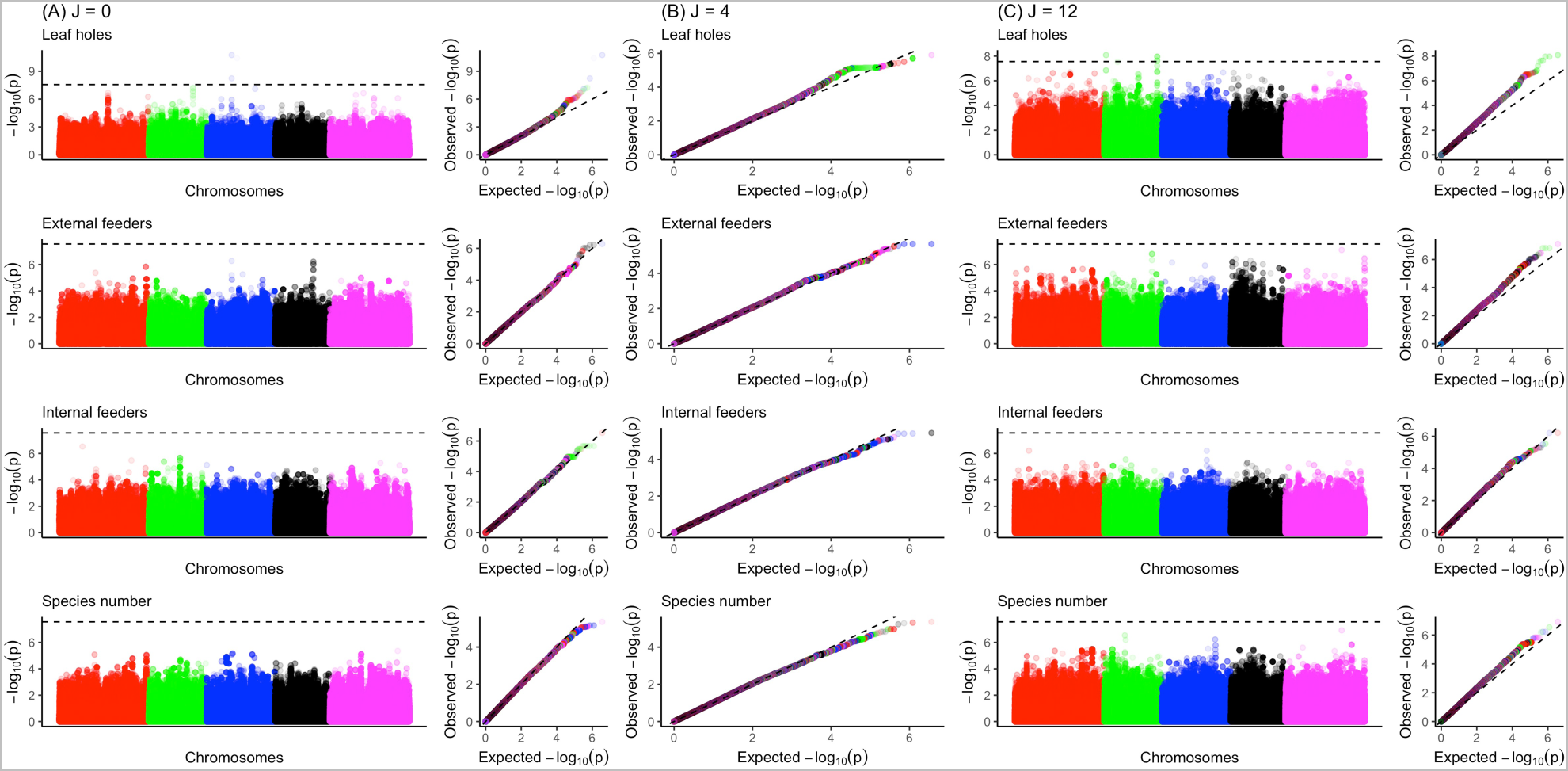
Manhattan and quantile-quantile (QQ) plots for the focal genotype effects and neighbor genotype effects on insect herbivory, abundance, and species number in Zurich. (A) *J* = 0 (Standard GWAS); (B) *J* = 4; and (C) *J* = 12. The horizontal dashed line of the Manhattan plot indicates a genome-wide Bonferroni threshold at *p* = 0.05. The gray dashed line of the QQ plots indicates the randomly expected *p*-values. The Manhattan plots at *J* = 4 are shown in the main Figure 2A.

**Figure S5.**
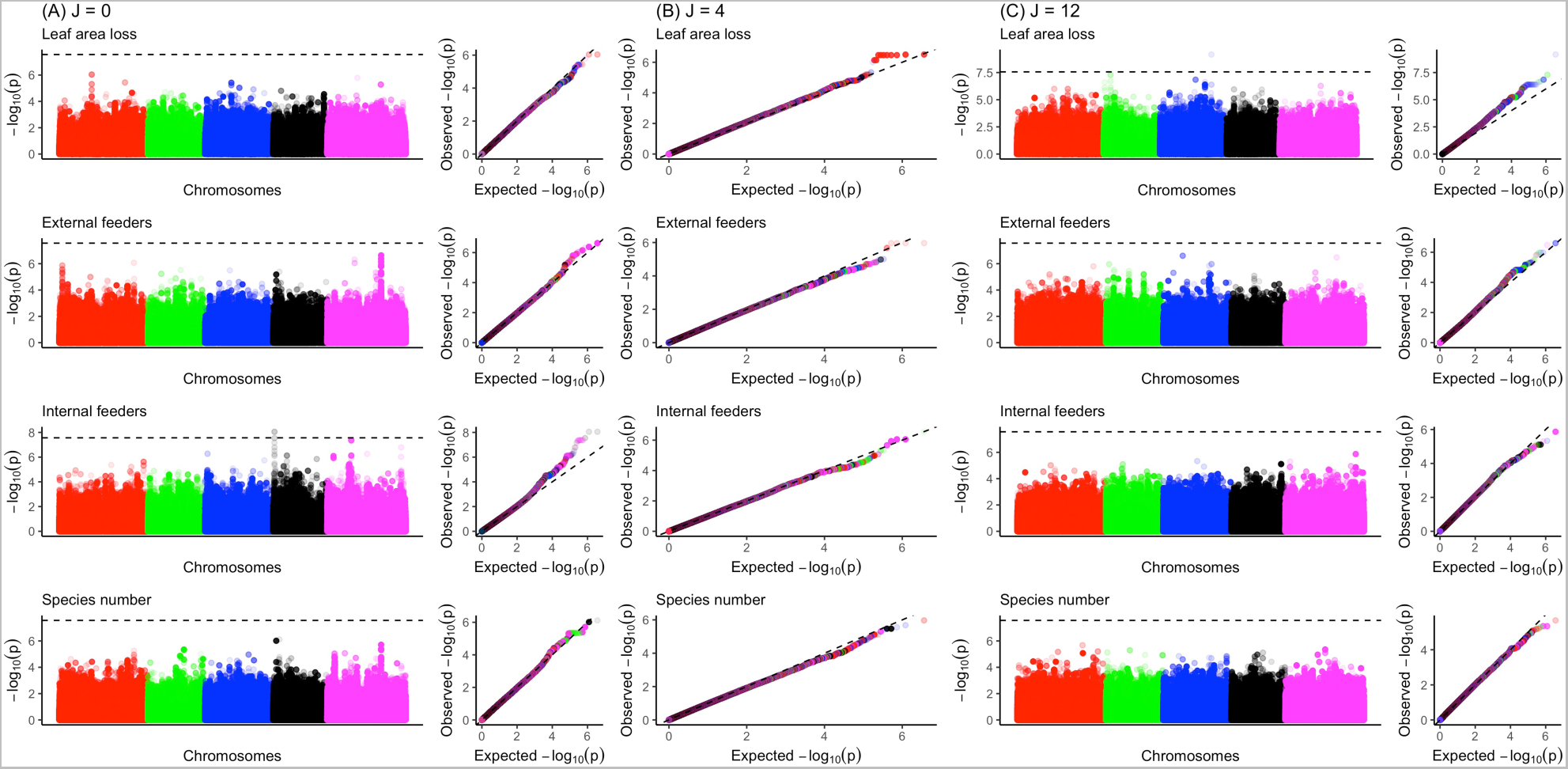
Manhattan and quantile-quantile (QQ) plots for the focal genotype effects and neighbor genotype effects on insect herbivory, abundance, and species number in Otsu. (A) *J* = 0 (Standard GWAS); (B) *J* = 4; and (C) *J* = 12. The Manhattan plots at *J* = 4 are shown in the main Figure 2B.

**Figure S6.**
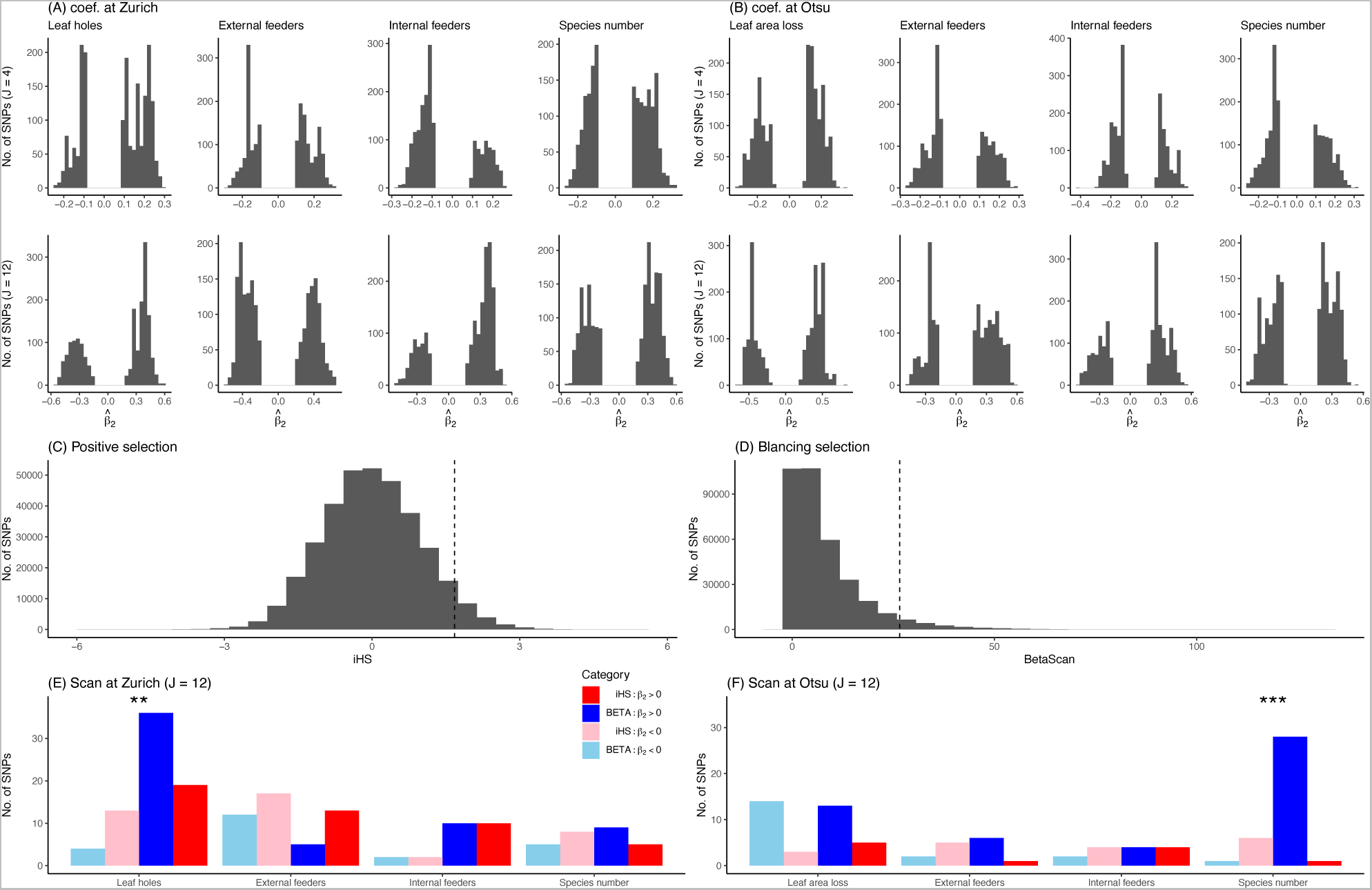
Comparison of top-scoring SNPs between association mapping and selection scans. (A and B) Distribution of the estimated *β*_2_ among the top 0.1%-scoring GWAS SNPs. (C and D) Genome-wide distribution of the indices of positive directional selection (iHS) and balancing selection (BetaScan) at MAF > 0.05. The vertical lines indicate 95 percentiles. (E and F) The number of SNPs shared between the selection scan (> top 5%) and the GWAS (> top 0.1%) at *J* = 12. The blue and red bars indicate balancing (BETA) and positive directional selection (iHS) indices with positive (darker color) or negative (paler color) 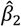, respectively. Asterisks indicate significant enrichment of the balancing selection between positive and negative 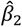; *** *p* < 0.001; ** *p* < 0.01 by Fisher tests.

**Figure S7.**
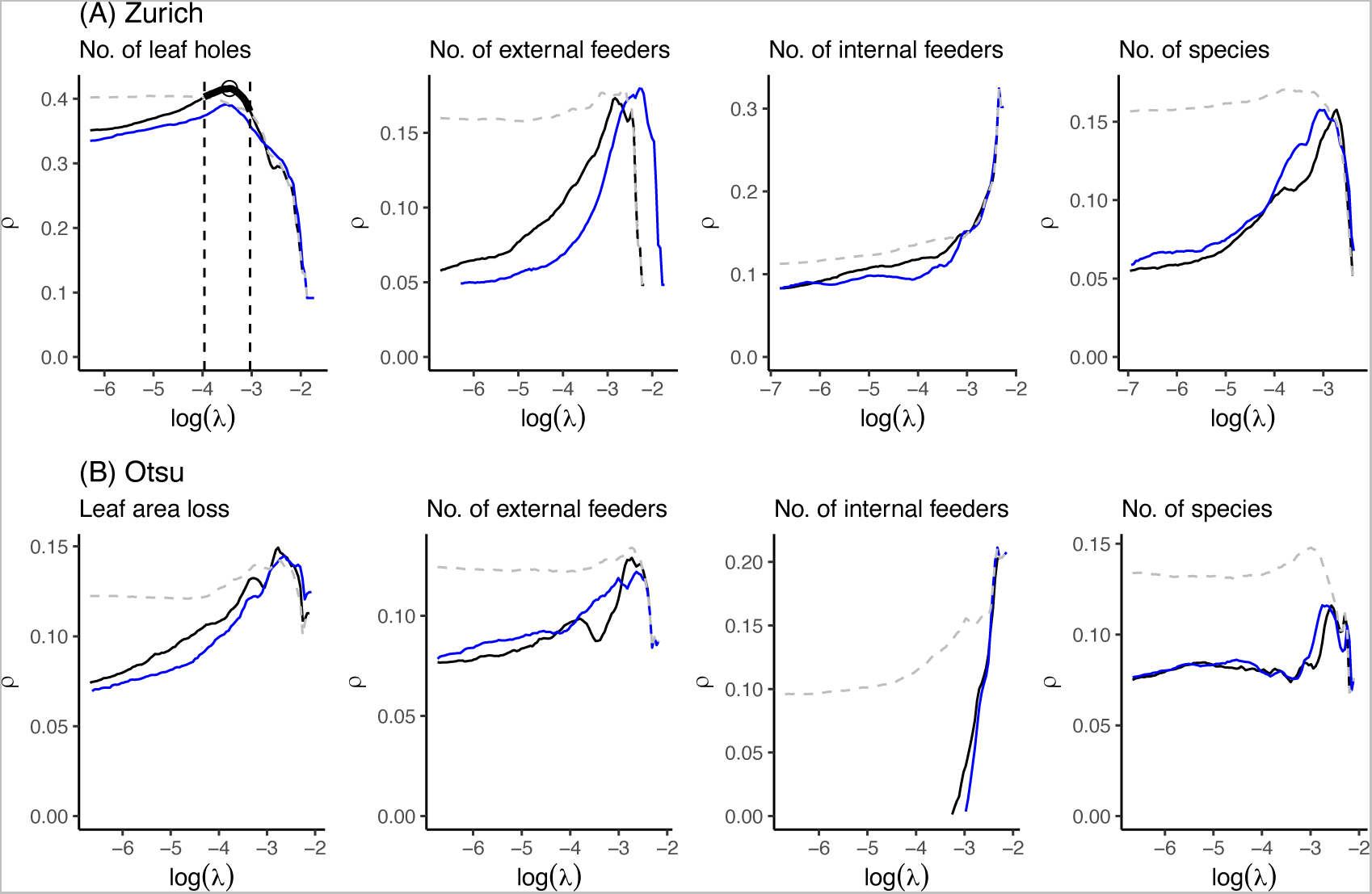
Rank correlations between observed and predicted phenotypes over a series of LASSO regularization parameters *λ* in (A) Zurich and (B) Otsu. Solid lines indicate the results of models including both focal and neighbor genotypes, while dashed lines indicate those without neighbor genotypes. The colors of the solid lines represent the reference space of the neighbor effects: *J* = 4 (black); *J* = 12 (blue). In Zurich (A), neighbor-including LASSO (*J* = 4; highlighted by a bold black line) achieved higher correlations with the number of leaf holes than neighbor-excluding LASSO (*J* = 0; gray dashed line) at more stringent regularization around the maximum *ρ* at ln(*λ*) = −3.1. An open circle highlights *λ* that yielded the maximum *ρ* = 0.416 for the neighbor-including LASSO at *J* = 4 in addition to *ρ* = 0.391 for the neighbor-excluding LASSO at *J* = 0 as mentioned in the main text. The two dashed vertical lines highlight the range of *λ* where the neighbor-including LASSO outperformed the neighbor-excluding LASSO, providing nonzero estimated SNPs in Figures S8A and B.

**Figure S8.**
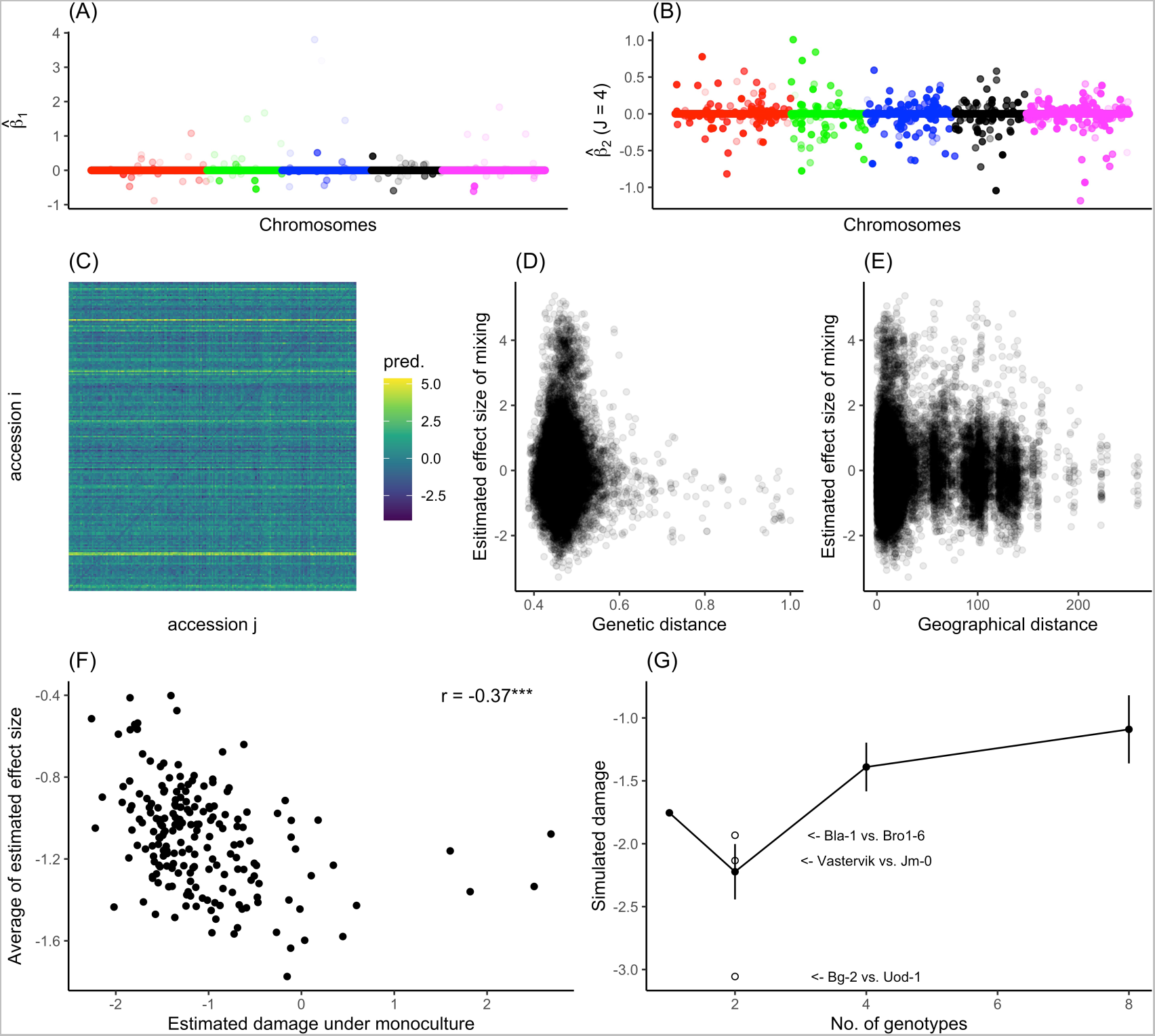
Selected SNPs and the estimated effects of mixed planting on leaf holes using LASSO. Focal (A) and neighbor (B) genotype effects estimated by LASSO for leaf holes at *J* = 4. (C) Heatmap showing the estimated damage between the accession *i* and *j*. Diagonal elements indicate the monoculture between the same accessions *i* and *i*, while the off-diagonal elements indicate the pairwise mixture between the accession *i* and *j*. (D and E) Pairwise effects plotted against the genetic distance (D) or geographical distance (E) between the two accessions. (F) Estimated herbivore damage under the virtual mixture condition plotted against that under monoculture conditions. A single plot indicates single accession. The y-axis shows the average of the estimated effect size (Fig. 3A) among 199 counterpart accessions for each focal accession on the x-axis. Pearson’s correction *r* and its level of significance from zero (*** *p* < 0.001) are also shown within a plot. This negative correction became larger (*r* = *–*0.424,*** *p* < 0.001) when four outlier accessions were omitted from the x-axis at > 1. (G) Predicted damage (mean ± SD) plotted against the number of randomly selected genotypes showing a positive effect size in Figure 3A.

**Figure S9.**
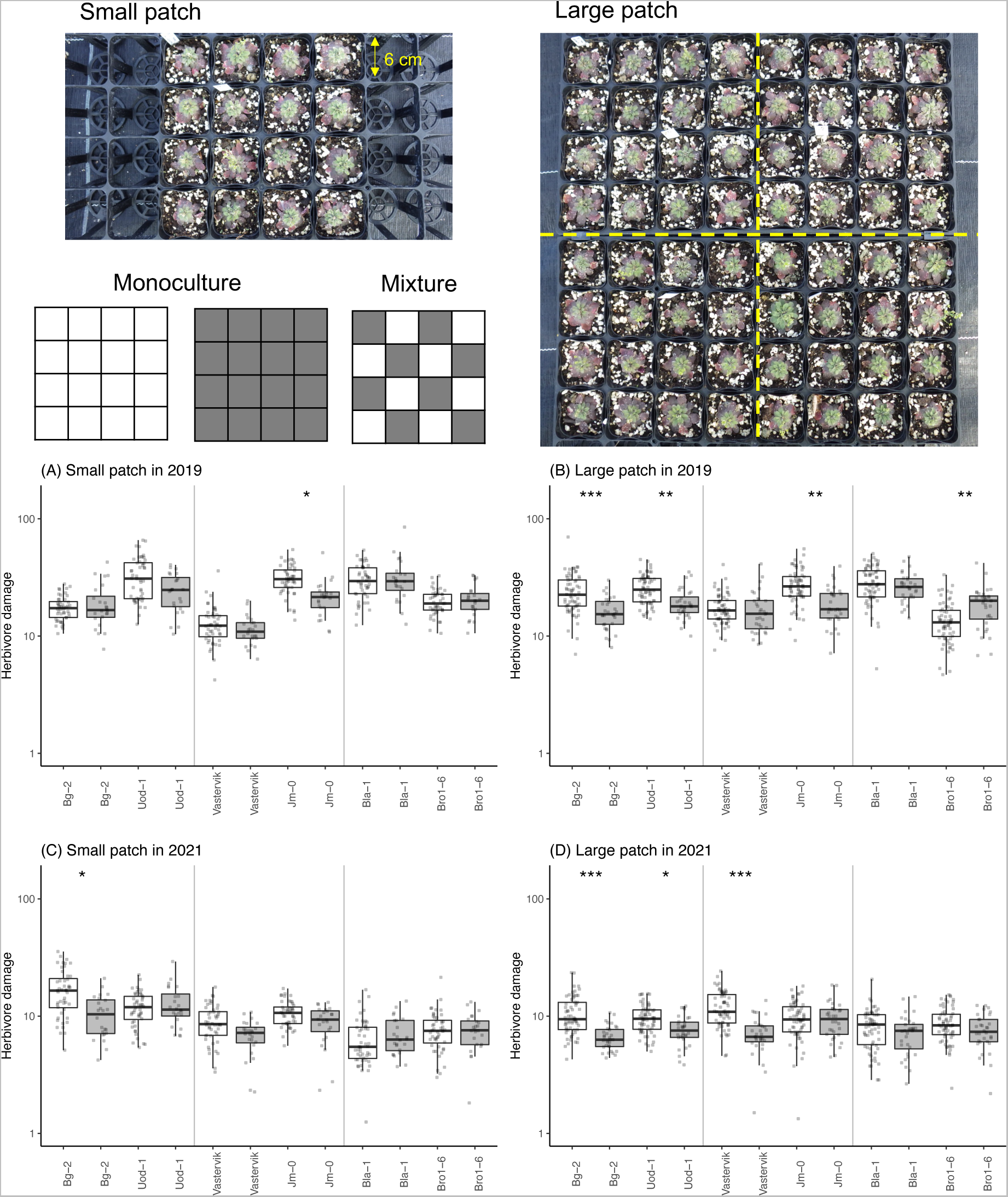
Experimental settings and the effects of mixed planting on herbivore damage in different years and patch conditions. The photographs display small and large patches. Yellow dashed lines in the large patches represent split subplots considered as a random effect in linear mixed models (see “Mixed planting experiment” in the Supplementary Materials and Methods). (A-D) Herbivore damage on the y-axis represents the number of leaf holes divided by the initial plant size (no./cm) on a logarithmic scale. White and gray boxes indicate the monoculture and mixture conditions, respectively. Asterisks indicate a significant difference in the estimated marginal means between the monoculture and mixture conditions: ** *p* < 0.01; *** *p* < 0.001.

**Figure S10.**
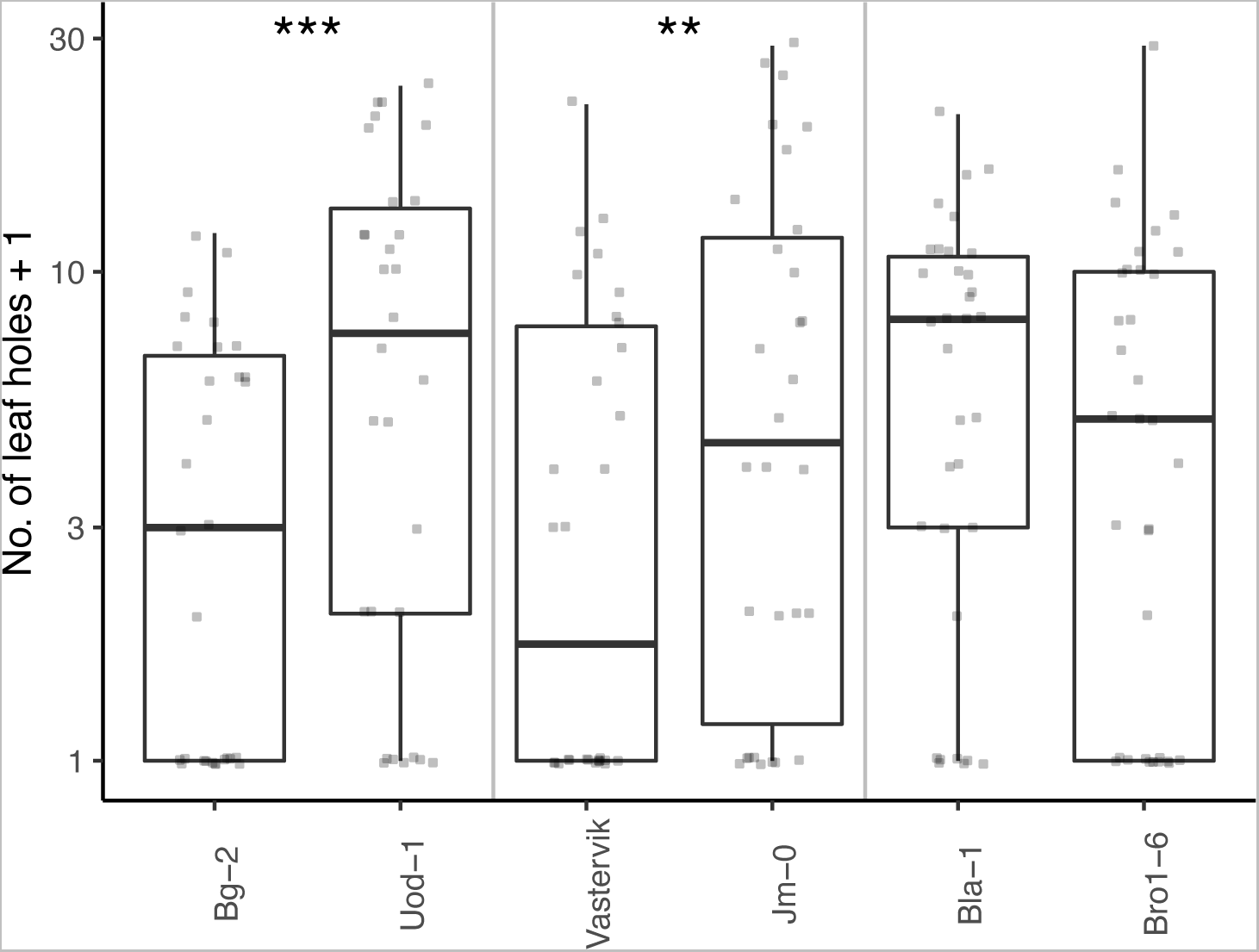
Paired choice experiments using adult flea beetles and three pairs of *Arabidopsis thaliana* accessions under laboratory conditions. Asterisks indicate significant differences between a pair of genotypes (Table S7): ** *p* < 0.01; *** *p* < 0.001.

**Table S1.** List of Arabidopsis thaliana accessions used in this study. (*see another file, TableS_AccessionList.csv*).

**Table S2.** List of arthropod species observed in this study. NA indicates ‘not applicable’. *Only this species is a non-insect arthropod. ^†^According to *mtCOI* sequences, the yellow-striped flea beetles in Zurich included two species, *P. striolata* and *P. undulata*, but they could not be identified by their appearance; therefore, these two species were counted as one morpho-species in this study.

| Common name | Scientific name | Order | Feeding habit | Host range | Pres./abs. Zurich | Pres./abs. Otsu | Abbrev. |
| --- | --- | --- | --- | --- | --- | --- | --- |
| Yellow-striped flea beetle | *Phyllotreta striolata*<sup>†</sup> | Coleoptera | Chewer | Oligophagous | + | + | Ps |
| Black flea beetle | *Phyllotreta astrachanica* | Coleoptera | Chewer | Oligophagous | + | - | Pa |
| Cabbage flea beetle | *Psylliodes punctifrons* | Coleoptera | Chewer | Oligophagous | - | + | Pp |
| Vegetable weevil | *Listroderes costirostris* | Coleoptera | Chewer | Polyphagous | - | + | Lc |
| Diamondback moth | *Plutella xylostella* | Lepidoptera | Chewer | Oligophagous | + | + | Px |
| Small cabbage white butterfly | *Pieris rapae* | Lepidoptera | Chewer | Oligophagous | + | + | Pr |
| Cabbage looper | *Trichoplusia ni* | Lepidoptera | Chewer | Polyphagous | + | + | Tn |
| Turnip sawfly | *Athalia rosae* | Hymenoptera | Chewer | Oligophagous | + | + | Ar |
| Garden springtail* | *Bourletiella hortensis* | Collembola | Chewer | Polyphagous | + | + | Bh |
| Cabbage bug | *Eurydema rugosa* | Hemiptera | Sucker | Oligophagous | - | + | Er |
| Green peach aphid | *Myzus persicae* | Hemiptera | Sucker | Polyphagous | + | + | Mp |
| Mustard aphids | *Lipaphis erysimi* | Hemiptera | Sucker | Oligophagous | + | + | Le |
| Cabbage aphids | *Brevicoryne brassicae* | Hemiptera | Sucker | Oligophagous | + | - | Bb |
| Flower thrip | *Frankliniella intonsa* | Thysanoptera | Sucker | Polyphagous | + | + | Fi |
| Western flower thrip | *Frankliniella occidentalis* | Thysanoptera | Sucker | Polyphagous | + | + | Fo |
| Diamondback moth | *Cotesia vestalis* | Hymenoptera | Carnivore | NA | + | + | Cv |

| *Common name* | *Scientific name* | *Order* | *Feeding habit* | *Host range* | *Pres./abs. Zurich* | *Pres./abs. Otsu* | *Abbrev.* |
| --- | --- | --- | --- | --- | --- | --- | --- |
| parasitoid |  |  |  |  |  |  |  |
| Larvae of hoverfly | Syrphinae sp. | Diptera | Carnivore | NA | + | + | Sy |
| Seven-spot ladybird | *Coccinella septempunctata* | Coleoptera | Carnivore | NA | + | + | Cs |
| (Parasitoid wasp indicated by mummified aphids) | NA | Hymenoptera | Carnivore | NA | + | + | mummy |

**Table S3.**
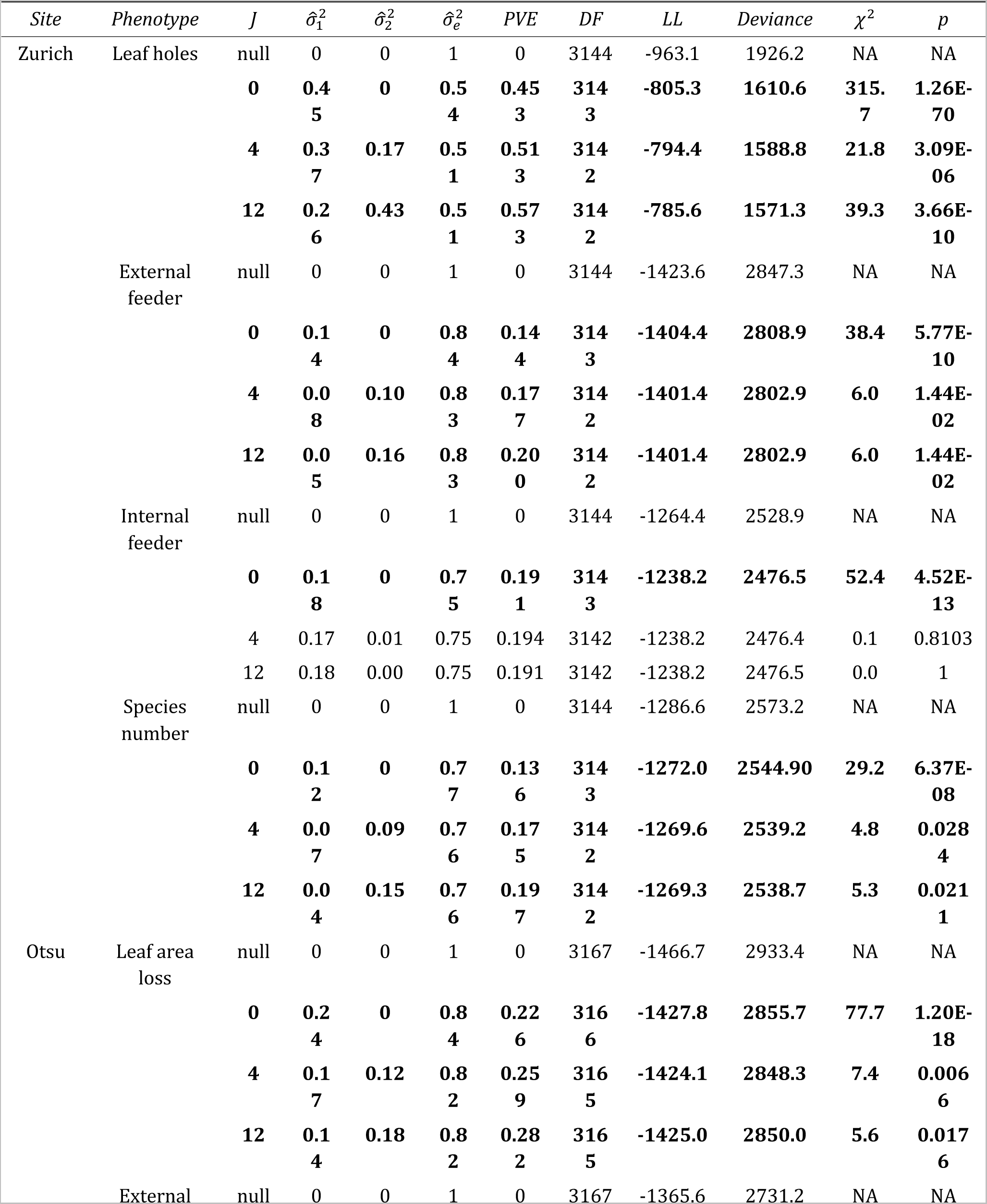
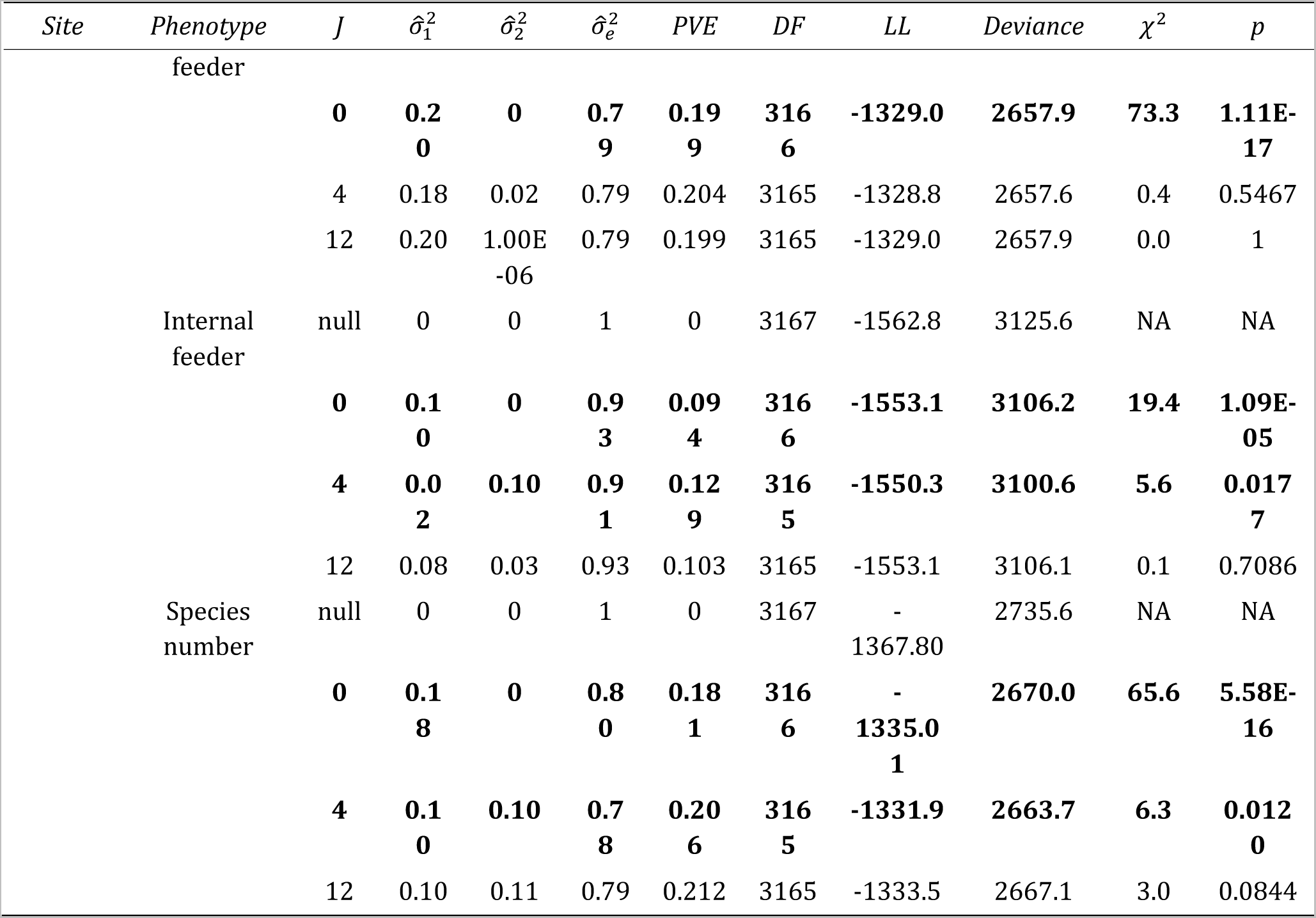
Likelihood ratio tests for variance component parameters in the standard and Neighbor GWAS. The deviance at *J* = 0 was tested against the null deviance, and the deviance at *J* = 4 and 12 was tested that at *J* = 0. Abbreviations: PVE, proportion of phenotypic variation explained by genetic factors; DF, degree of freedom; LL, log-likelihood. Bold values highlight *p* < 0.05.

**Table S4.** List of candidate genes from GWAS of insect herbivory, abundance, and species number in the Zurich and Otsu site. The possibility of positive or balancing selection was also annotated to each SNP (*see another file, TableS_GWAScandidate.xlsx*).

**Table S5.** Linear mixed models for analyzing the effects of mixed planting on leaf holes. (A) Analysis of variance (ANOVA) comparing models with or without a single explanatory variable. (B) Estimated marginal means of the effects of the mixture conditions in the full model. The positive estimates indicated an increase in the number of leaf holes in the monoculture relative to the mixture conditions. Bold values highlight *p* < 0.05. Abbreviations: DF: degree of freedom; SE: standard error.

| *Explanatory variable* | *Sum of squared* | *Mean squared* | *Numerator DF* | *Denominator DF* | *F* | *p* |
| --- | --- | --- | --- | --- | --- | --- |
| **Study year** | **44.022** | **44.022** | **1** | **111.05** | **396.190** | **2.20E-16** |
| Patch size | 0.172 | 0.172 | 1 | 111.06 | 1.551 | 0.215626 |
| **Mixture condition** | **1.208** | **1.208** | **1** | **111.06** | **10.876** | **0.001309** |
| **Genotype** | **7.765** | **1.553** | **5** | **142.78** | **13.977** | **4.06E-11** |
| Patch × Mix | 0.062 | 0.062 | 1 | 111.06 | 0.561 | 0.455632 |
| **Patch × Geno** | **1.675** | **0.335** | **5** | **276.2** | **3.015** | **0.011464** |
| Mix × Geno | 1.053 | 0.211 | 5 | 141.38 | 1.895 | 0.098884 |

| *Genotypes* | *Estimate* | *SE* | *DF* | *t-value* | *adjusted p* |
| --- | --- | --- | --- | --- | --- |
| **Bg-2** | **0.307** | **0.093** | **125** | **3.30** | **0.0013** |
| **Uod-1** | **0.188** | **0.093** | **125** | **2.02** | **0.0455** |
| **Vastervik** | **0.246** | **0.093** | **125** | **2.64** | **0.0093** |
| **Jm-0** | **0.207** | **0.093** | **125** | **2.22** | **0.0282** |
| Bla-1 | -0.058 | 0.093 | 125 | -0.62 | 0.536 |
| Bro1-6 | 0.011 | 0.093 | 125 | 0.12 | 0.905 |

**Table S6.** Linear mixed models for analyzing the effects of mixed planting on leaf holes under different patch conditions over two years. As shown in Table S5, the upper and lower tables of each panel display the results of the analysis of variance and estimated marginal means, respectively. Bold values highlight *p* < 0.05.

| *Explanatory variable* | *Sum of squared* | *Mean squared* | *Numerator DF* | *Denominator DF* | *F* | *p* |
| --- | --- | --- | --- | --- | --- | --- |
| Mixture condition | 0.2535 | 0.25348 | 1 | 18.025 | 2.6619 | 0.1201 |
| **Genotype** | **15.4601** | **3.09201** | **5** | **26.911** | **32.4703** | **1.37E-10** |
| Mix × Geno | 0.9509 | 0.19017 | 5 | 26.911 | 1.9971 | 0.1113 |

| *Genotypes* | *Estimate* | *SE* | *DF* | *t-value* | *adjusted p* |
| --- | --- | --- | --- | --- | --- |
| Bg-2 | -0.038 | 0.136 | 22.6 | -0.28 | 0.783 |
| Uod-1 | 0.277 | 0.136 | 22.6 | 2.04 | 0.053 |
| Vastervik | 0.093 | 0.136 | 22.6 | 0.69 | 0.500 |
| **Jm-0** | **0.366** | **0.136** | **22.6** | **2.70** | **0.013** |
| Bla-1 | -0.028 | 0.136 | 22.6 | -0.21 | 0.838 |
| Bro1-6 | -0.042 | 0.136 | 23.1 | -0.31 | 0.761 |

(B) Large patch in 2019
| *Explanatory variable* | *Sum of squared* | *Mean squared* | *Numerator DF* | *Denominator DF* | *F* | *p* |
| --- | --- | --- | --- | --- | --- | --- |
| **Mixture condition** | **1.0855** | **1.08555** | **1** | **27.114** | **8.8754** | **0.006031** |
| **Genotype** | **14.049** | **2.80979** | **5** | **51.573** | **22.9728** | **4.71E-12** |
| **Mix × Geno** | **3.9201** | **0.78403** | **5** | **51.573** | **6.4102** | **0.000106** |

| *Genotypes* | *Estimate* | *SE* | *DF* | *t-value* | *adjusted p* |
| --- | --- | --- | --- | --- | --- |
| **Bg-2** | **0.3653** | **0.0951** | **43.2** | **3.84** | **0.0004** |
| **Uod-1** | **0.2873** | **0.0951** | **43.2** | **3.02** | **0.0042** |
| Vastervik | 0.0544 | 0.0951 | 43.2 | 0.572 | 0.5701 |
| **Jm-0** | **0.2952** | **0.0951** | **43.2** | **3.103** | **0.0034** |
| Bla-1 | 0.0508 | 0.096 | 44.6 | 0.53 | 0.5988 |
| **Bro1-6** | **-0.2971** | **0.0951** | **43.2** | **-3.123** | **0.0032** |

(C) Small patch in 2021
| *Explanatory variable* | *Sum of squared* | *Mean squared* | *Numerator DF* | *Denominator DF* | *F* | *p* |
| --- | --- | --- | --- | --- | --- | --- |
| Mixture condition | 0.16583 | 0.16583 | 1 | 18 | 1.6168 | 0.219727 |
| **Genotype** | **2.70265** | **0.54053** | **5** | **22.208** | **5.2699** | **0.002441** |
| Mix × Geno | 0.76187 | 0.15237 | 5 | 22.208 | 1.4856 | 0.234422 |

| *Genotypes* | *Estimate* | *SE* | *DF* | *t-value* | *adjusted p* |
| --- | --- | --- | --- | --- | --- |
| **Bg-2** | **0.4537** | **0.207** | **19.9** | **2.188** | **0.0408** |
| Uod-1 | -0.0631 | 0.207 | 19.9 | -0.304 | 0.7642 |
| Vastervik | 0.273 | 0.207 | 19.9 | 1.316 | 0.203 |
| Jm-0 | 0.2121 | 0.207 | 19.9 | 1.023 | 0.3188 |
| Bla-1 | -0.1145 | 0.207 | 19.9 | -0.552 | 0.5869 |
| Bro1-6 | 0.0101 | 0.207 | 19.9 | 0.049 | 0.9618 |

(D) Large patch in 2021
| *Explanatory variable* | *Sum of squared* | *Mean squared* | *Numerator DF* | *Denominator DF* | *F* | *p* |
| --- | --- | --- | --- | --- | --- | --- |
| **Mixture condition** | **2.6127** | **2.61273** | **1** | **27** | **23.2567** | **4.91E-05** |
| Genotype | 1.0728 | 0.21455 | 5 | 44.912 | 1.9098 | 0.111524 |
| **Mix × Geno** | **2.4543** | **0.49086** | **5** | **44.912** | **4.3693** | **0.002517** |

| *Genotypes* | *Estimate* | *SE* | *DF* | *t-value* | *adjusted p* |
| --- | --- | --- | --- | --- | --- |
| **Bg-2** | **0.4157** | **0.109** | **37.2** | **3.82** | **0.0005** |
| **Uod-1** | **0.2275** | **0.109** | **37.2** | **2.091** | **0.0435** |
| **Vastervik** | **0.5491** | **0.109** | **37.2** | **5.046** | **<.0001** |
| Jm-0 | 0.0121 | 0.109 | 37.2 | 0.111 | 0.9121 |
| Bla-1 | 0.1089 | 0.109 | 37.2 | 1 | 0.3236 |
| Bro1-6 | 0.1398 | 0.109 | 37.2 | 1.284 | 0.207 |

**Table S7.** Generalized linear models analyzing the number of leaf holes in paired choice experiments. Likelihood ratio tests based on the deviance and *χ*^2^-distribution are shown for the three pairs of genotypes. Bold values highlight *p* < 0.05.

| *Explanatory variable* | *DF* | *Deviance* | *p* |
| --- | --- | --- | --- |
| **Arena** | **14** | **25.42** | **0.031** |
| **Genotype** | **1** | **13.35** | **0.0003** |
| Residual | 44 | 69.36 | — |

(B) Vastervik vs. Jm-0
| *Explanatory variable* | *DF* | *Deviance* | *p* |
| --- | --- | --- | --- |
| Arena | 14 | 20.89 | 0.105 |
| **Genotype** | **1** | **5.71** | **0.017** |
| Residual | 44 | 64.57 | — |

| *Explanatory variable* | *DF* | *Deviance* | *p* |
| --- | --- | --- | --- |
| Arena | 16 | 24.90 | 0.072 |
| Genotype | 1 | 0.87 | 0.352 |
| Residual | 50 | 81.97 | — |

**Table S8.** List of candidate genes from LASSO of herbivore damage in the Zurich site. Estimated focal and neighbor genotype effects *β*_1_ and *β*_2_ are shown for non-zero SNPs (*see another file, TableS_LASSOcandidate_HolesZurich.xlsx*).

**Table S9.** Gene ontology (GO) enrichment analyses for candidate genes from LASSO. The GO terms were reduced using the REVIGO algorithm. The tab (A) and (B) show the list of GO terms for candidate genes from positive and positive *β*_2_, respectively (*see another file, TableS_LASSO_REVIGO.xlsx*).

